# The interplay of facilitation and competition drives the emergence of multistability in dryland plant communities

**DOI:** 10.1101/2024.01.09.573035

**Authors:** Benôıt Pichon, Isabelle Gounand, Sophie Donnet, Sonia Kéfi

**Affiliations:** ISEM, Univ Montpellier, CNRS, EPHE, IRD, F-34095 Montpellier, France; Institut d’écologie et des sciences de l’environnement (iEES Paris), Sorbonne Université, CNRS, UPEC, CNRS, IRD, INRA, 75005 Paris, France; UMR MIA-Paris, AgroParisTech, INRA, Université Paris-Saclay, 75005, Paris, France; Santa Fe Institute, 1399 Hyde Park Road, Santa Fe, NM 87501, USA

**Keywords:** Drylands, multistability, alternative stable states, feedbacks, spatial model, plant community, niche construction, facilitation, spatial scale, desertification

## Abstract

Species are wrapped in a set of feedbacks within communities and with their abiotic environment, which can can generate alternative stable states. So far, research on alternative stable states has mostly focused on systems with a small number of species and a limited diversity of interaction types. Here, we analyze a spatial model of plant community dynamics in drylands, where each species is characterized by a strategy, and interact through facilitation and competition. Our work identifies three different types of multistability emerging from the interplay of competition and facilitation. Under low-stress levels, the community organizes in small groups of coexisting species maintained by space and facilitation, while under higher stress levels, positive feedbacks from competition and facilitation lead to the dominance of a single species before desertification happens. Our study paves the way for bridging community ecology and alternative states theory in a common framework.

## Introduction

Due to the inherent complexity of ecological systems, species, and their environment are involved in complex sets of feedbacks (DeAngelis et al., 1986; Odling-Smee et al., 1996). Feedback loops are a set of interactions from a given species that cycle, through the rest of the interaction network, and goes back to the focal species (Pichon et al., 2023). Their sign can be positive, in which case perturbations are amplified through this loop, or negative, in which case perturbations are dampened. When positive feedbacks are strong enough, small changes in environmental conditions can trigger abrupt shifts between alternative states. This can happen when multiple ecosystem states are possible for a given set of environmental conditions (also referred to as ”multistability”; Scheffer et al., 2001; Kéfi et al., 2016a).

Such abrupt transition between alternative states have been shown in many systems such as shallow lakes and savannas (Scheffer, 2009), but also climate (Rial et al., 2004; Lenton et al., 2008) and past civilization demography (Janssen et al., 2003; Centeno et al., 2023). A well-documented example comes from drylands, where plants improve their local environment, increase the local availability of water and the cycling of nutrients via their rooting system, and limit water evaporation by shading, which ultimately benefit plant recruitment, growth, and survival (*i.e.,* positive niche construction; Callaway, 2007; Odling-Smee et al., 2013; Filazzola and Lortie, 2014). With increasing aridity, vegetation cover decreases in drylands because of lower seed recruitment, therefore limiting the positive feedback generated by facilitation in the ecosystem, which in turn negatively affects vegetation cover. Eventually, facilitation can no longer maintain the vegetation, and desertification occurs with an abrupt loss of cover (Schlesinger et al., 1990; Rietkerk and van de Koppel, 1997; Kéfi et al., 2007). So far, while many theoretical studies have investigated the emergence of alternative states in drylands, most of them have relied on one or two species (*e.g.,* Rietkerk and van de Koppel, 1997; Gilad et al., 2007; Danet et al., 2021), thereby leaving the question of the role of species diversity largely open (Kéfi et al., 2022).

In other ecological systems, studies have investigated how alternative stable states emerge in species rich communities but with a single interaction type such as trophic (De Roos and Persson, 2002; Downing et al., 2012; Karatayev et al., 2023), competitive (Gilpin and Case, 1976; van Nes and Scheffer, 2004) or mutualistic communities (Lever et al., 2014). These previous studies have unveiled that positive feedbacks emerging from density-dependent foraging or from competition between species could lead to alternative stable states between communities of different species compositions. However despite a rich theoretical background on communities characterized by either trophic, competitive or facilitative links, we still lack an understanding of the role of diverse interaction types in generating alternative community states.

In fact, in all ecosystems, species interact both positively and negatively (Fontaine et al., 2011; Pocock et al., 2012; Kéfi et al., 2016b). This creates interdependences between species, which can serve as a medium to propagate local perturbations to the whole community. Integrating this diversity of interactions can remarkably change our predictions of how species coexist (*e.g.,* Gross, 2008; Aubier, 2020; Kéfi et al., 2016b). For instance, the inclusion of positive interactions in competitive communities can increase the number of coexisting species (Kéfi et al., 2012; Gross, 2008) or the stability of communities facing small perturbations (Mougi and Kondoh, 2012). So far, in this emerging literature aiming at integrating the diversity of interaction types into ecological theory, multistability has been largely overlooked. Recently, Lever et al. (2020) showed that strong mutualism in a model of plant-pollinator communities can generate abrupt species collapse, while strong competition between pollinators generates transitions from stable to oscillatory dynamics. In drylands, a combination of facilitation and competition between plants generate spatial patterns and determine the dynamics and stability of the communities (Rietkerk, 2004). In addition to species interactions, dispersal is a key trait for recruitment since seeds need to access fertile areas that are often located in the neighborhood of facilitating species (Aguiar et al., 1994; Flores and Jurado, 2003; Pueyo et al., 2008). When combined together, these mechanisms allow dryland sites to host multiple species (typically between 5 and 35; Maestre et al., 2012) with a range of strategies, from competitivity for water to stress-tolerance (Valiente-Banuet et al., 2006; Angert et al., 2009; Graff and Aguiar, 2017). Acknowledging this diversity of strategies and of interaction types, rather than seeing drylands through the lens of a single species, may change our understanding of the stability of these ecosystems. In particular, with the ongoing increase in levels of stress imposed on dryland species due to climate change and anthropogenic pressures (Burrell et al., 2020), we expect dryland plant communities to respond differently depending on their functional composition, and the interplay between competition and facilitation between plant species (Ruppert et al., 2015; Nunes et al., 2017).

Extending multistability in a spatial context in drylands, we here investigate the emergence of multistability in dryland plant communities using an individual-based model of plant community dynamics, in which each plant is characterized by its ecological strategy (tolerance to stress *versus* competitivity for water) and interacts through facilitation and competition with others. We explore how the emergence of multistability is modulated by the strength of facilitation and competition as well as by the dispersal strategies of the plants. Our results reveal three types of multistability, all emerging from positive feedback loops but driven by different mechanisms.

## Methods

### Model description

We propose a model which expands a single-species model of vegetation in dryland ecosystems (Kéfi et al., 2007) to *N* interacting species (Fig. 1). The landscape is represented by a grid of *n × n* sites, each measuring approximately a square meter. Each site is in one of the following states: fertile soil (0, colonizable by vegetation), degraded (*−*, not colonizable), or vegetated (+*_i_*, *i.e.,* occupied by a plant species *i*, *i* = 1, . . . , *N*).

**Figure 1:**
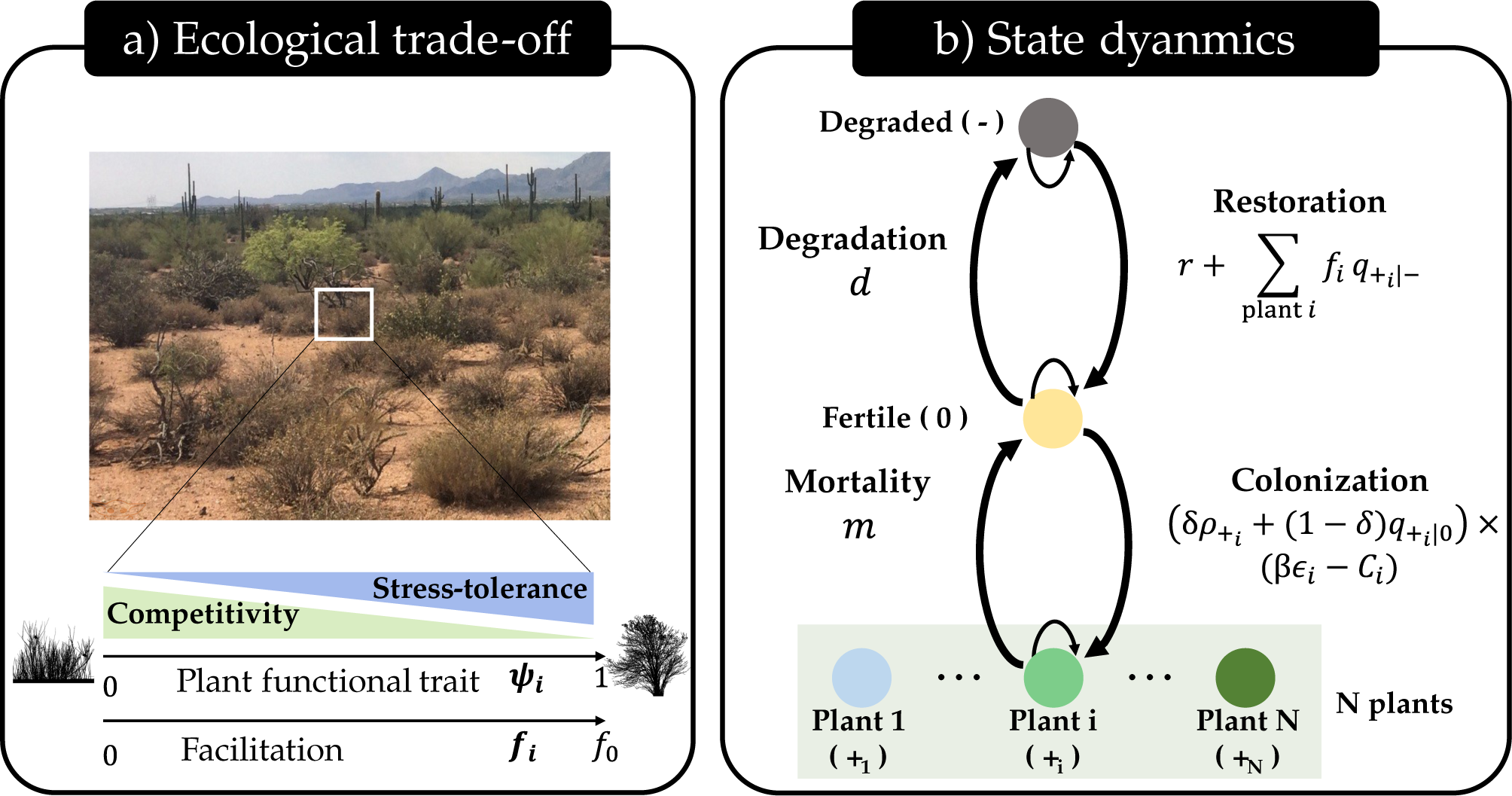
A plant community model for arid ecosystems. (a) We modelled the dynamics of vegetation in arid ecosystems where different functional strategies can coexist. Each species is characterized by a strategy that is its position (*ψ_i_*) along a linear trade-off between competitivity and stress-tolerance. A plant’s functional strategy determines its tolerance to abiotic stress, the intensity at which it facilitates the local environment, and its tolerance to water competition. (b) The model describes the dynamics of *N* + 2 states: *N* plants indexed by *i* (+*_i_*, *i* = 1, . . . , *N* ), the fertile soil (0) and degraded (*−*) sites, in blue/green, yellow, and grey, respectively. The functional strategy of plants determines the transition rates between each state. Symbols of transition rates are used in the model equations: degradation (*d*) and basal restoration (*r*) of a fertile site, plant-specific facilitation intensity ( *f_i_*), the local density of plants *i* surrounding a degraded site (*q*_+*i*_ *_|−_*; 4 nearest neighbors), global (*δρ*_+*i*_ ) and local (1 *− δ*)*q*_+*i |*0_) dispersal terms, production (*β*) and germination of seed (*ɛ_i_*), species-specific stress (*S_i_*) and competition (*C_i_*) experienced, and plant mortality (*m*). An illustration of the possible transitions can be found in Fig. S1.1.

#### Plant strategies

In arid (Liancourt et al., 2005; Angert et al., 2009; Graff and Aguiar, 2017), alpine (Choler et al., 2001) and salted ecosystems (He et al., 2012; Zhang and Tielbörger, 2019), species have been shown to be distributed along an ecological trade-off: some species have a strong competitive ability, while others are more tolerant to abiotic stress (Díaz et al., 2016). We therefore characterized each plant species with a strategy corresponding to a position, *ψ_i_*, along the competitivity-tolerance trade-off axis (*ψ_i_ ∈* [0, 1]; Fig. 1a). We assumed the trade-off of strategies to be linear (but see Appendix S2 for varying shapes). When *ψ_i_* = 1, species are stress-tolerant, but have a low competitivity against other plants when competing for resources (*i.e.,* high sensitivity to interspecific competition). At the other end of the trade-off (*ψ_i_* = 0), species are less sensitive to competition at the expanse of a sensitivity to abiotic stress (Butterfield and Briggs, 2011). In addition, following (Armas and Pugnaire, 2005; Liancourt et al., 2005; Fagundes et al., 2022), we assumed that contrary to competitive strategies, stress-tolerant ones are enhancing local conditions in their surrounding by increasing the regeneration rate of degraded sites into fertile ones (positive niche construction).

#### State transitions and spatial dynamics

We model the annual dynamics of each site in the landscape that change accordingly to four possible transitions: a fertile site can be colonized by a plant *i* with a probability *P*(0 *→* +*_i_*) or degraded by erosion at a rate *P*(0 *→ −*) (Fig. 1b). In addition, surrounding vegetation and spontaneous regeneration of soil create new fertile sites with a probability *P*(*− →* 0). Finally, plants can die, for instance by natural mortality or herbivory, with a probability *P*(+*_i_ →* 0) (Fig. S1.1 for illustration). From all the sites, we derive aggregated variables of the landscape that are the global density of each state across the landscape (*ρ_k_*, *k ∈ {−*, 0, +_1_, . . . , +*_N_}*), and the local density of state *k^′^* in the neighborhood (four-nearest neighbors) of a given site in state *k* (*q_k′ |k_*).

The recruitment of plant *i* on a given fertile site occurs with a probability:

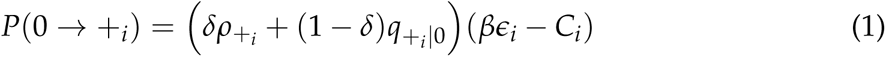

where *δ* (*resp.,* 1 *− δ*) is the fraction of seeds globally (*resp.,* locally) dispersed across the landscape (*resp.* in the neighborhood), *ρ*_+_*_i_* is the global density of plants *i*, *q*_+_*_i |_*_0_ is the local density of plant *i* near the fertile site, *β* is the number of seeds produced, *ɛ_i_* the germination rate of plant *i* in a fertile site and *C_i_* the competition experienced by plant *i* during recruitment. We further assumed that the probability of germination decreases linearly with the stress experienced by each plant *i* following: *ɛ_i_* = 1 *− S_i_*. Therefore, independently of their strategies, plants have an equal maximal germination rate of 1. In addition, because plants have different stress-tolerance abilities, the stress experienced by each plant *i*, *S_i_*, will depend on its ecological strategy:

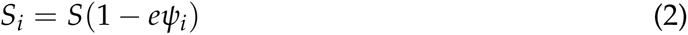

where *S* is the global abiotic stress imposed on the landscape and *e* is the maximal fraction of stress relaxed by stress-tolerant species. The larger *ψ_i_*, the lower stress experienced by plants, and the higher the germination rate.

Besides, on top of the spatial competition for the recruitment on fertile sites, we assumed a competition term (*C_i_*) for water driven by the global species density due to the redistribution of water across the landscape:

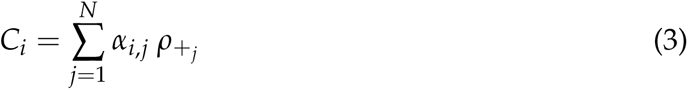

where *α_i_*_,*j*_ is the competition coefficient transcribing the negative effect of species *j* on species *i*. We set intraspecific interactions (*α_i_*_,*i*_) to *α*_0_, and assumed interspecific interaction between species to be modulated by three factors: *(i)* a basal level of competition (*α_e_*), *(ii)* the similarity in species strategies (*|ψ_i_ − ψ_j_|*), as functionally similar species tend to be more in competition (MacArthur and Levins, 1967; Cahill et al., 2008; Burns and Strauss, 2011), and *(iii)* the sensitivity of plant *i* to competition, which was approximated by *ψ_i_*:

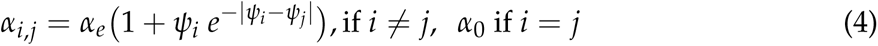

Therefore a competitive species (low *ψ*) is less sensitive to the competition experienced by all other species. Note that this shape of the competition function favors species at the edges of the trade-off because they have fewer competitors with similar strategies (Fig. S1.2).

The regeneration rate of degraded sites depends on the spontaneous regeneration rate of degraded site (*e.g.,* by rainfall events), and on the local density of established plants in the neighborhood of a degraded site *q*_+_*_i |−_*:

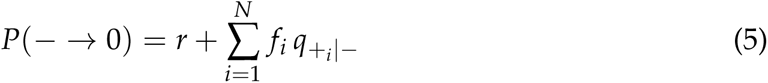

Stress-tolerant species are assumed to be higher facilitators compared to competitive ones: *f_j_*= *ψ_j_ f*_0_, where *f*_0_ is the maximal facilitation strength.

Plant mortality (*P*(+*_i_ →* 0)) occurs at rate *m* independent of the plant strategy (as discussed in Appendix S3), and turns a site occupied by vegetation to a fertile site. Finally, these fertile sites are lost by natural erosion at a rate of *P*(0 *→ −*) = *d*. In sum, equations (1)-(5) define the spatio-temporal dynamics of the plant community:

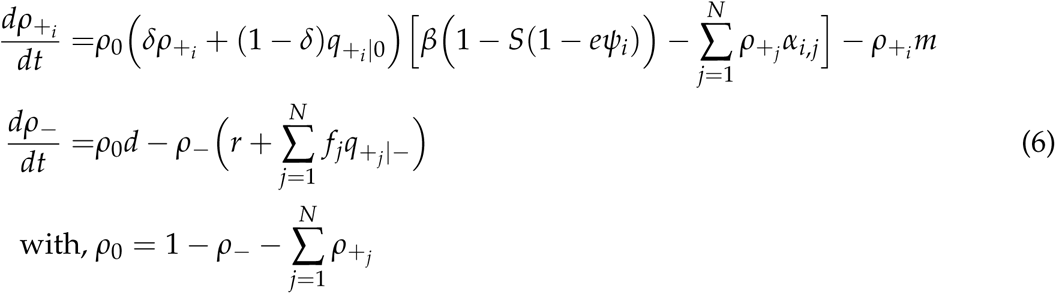

We parameterized the model following Kéfi et al. (2007) (see Appendix S3 for details).

### Analyzing the emergence of multistability

#### Approximating the stochastic spatial model

We approached the dynamics of our spatially explicit model using two deterministic approximations. First, a mean-field model that ignores the spatial structure and only follows the global densities of each state (*ρ*_+_*_i_* ; Appendix S4). Secondly, we used the pair-approximation model that follows global densities but also the pair densities (*e.g.,* pair (+*_i_*, 0); see Appendix S5). Comparing the results between the two approximations allows to understand the effect of space on the dynamics of the system. Because the PA model gives quantitatively similar results as the stochastic spatial model (Fig. S1.3), we used the PA model for all analyses in the main text and illustrated some dynamics using stochastic simulations.

#### Building the bifurcation diagrams

To detect multistability, we computed the cover of each species along the abiotic stress gradient (*S*) for multiple different initial conditions. Because some community dynamics may result in the same total vegetation cover despite different species compositions, we used a community index metric that characterize each different alternative state with a single number (as detailed in Appendix S3).

#### Mechanistic understanding with two species

To gain a mechanistic understanding of the model behavior, we start our analysis with a two-species community with two extreme strategies: a species being stress-tolerant (*ψ* = 1), the other being competitive (*ψ* = 0). We tracked the emergence of multistability by varying the interspecific competition strength (*α_e_*), and the dispersal of seeds.

We considered a local dispersal scenario as it is a common adaptation of plants in drylands (*i.e.,* proxichory with *δ* = 0.1; Fllner and Shmida, 1981) and a global dispersal scenario (*δ* = 0.9, mediated by wind or herbivores).

#### Multistability in species-rich communities

We scaled up to species-rich communities to understand the different types of multistability that may emerge. We worked on simulations with *N ∈ {*5, 15, 25*}* species. For each community richness, species strategies were equally split along the trade-off axis such that the dissimilarity of strategy between each consecutive pair of species *i* and *i* + 1, *|ψ_i_ − ψ_i_*_+1_*|*, was constant. We tracked the emergence of multistability by computing 250 bifurcation diagrams with random initial species cover while keeping their sum to 0.8. All simulations were made with Julia (1.7.3) using the *DifferentialEquation* package and analyzed with R (4.1.0).

## Results

### Community succession

In two-species simulations starting with individuals from a stress-tolerant and a competitive species randomly spread in the landscape, for a low-stress level and a weak interspecific competition, the two species can coexist (Fig. 2a yellow area).

**Figure 2.**
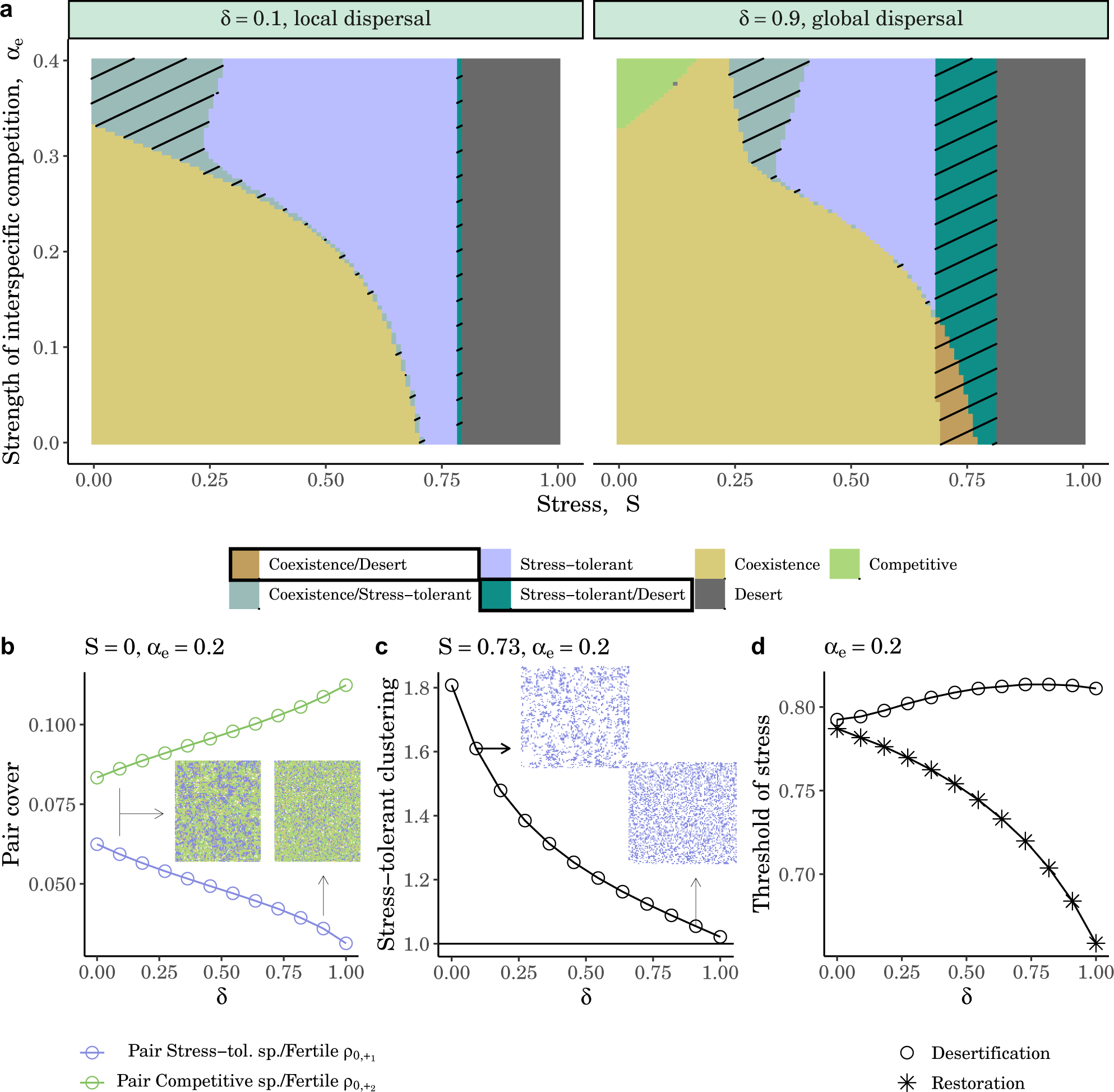
Dispersal scale and competition strength modulate the emergence of bistability. The community is composed of two species, one being stress-tolerant (*ψ*_1_ = 1) and the other a competitive (*ψ*_2_ = 0). (a) State diagram of the different community states along a gradient of abiotic stress (*S*) and interspecific competition strength (*α_e_*). The community state is displayed for two values of dispersal (in columns): local (*δ* = 0.1, *i.e.* 90% of the seeds are dispersed in the local neighborhood of the plant) and global dispersal (*δ* = 0.9). Areas of bistability have been hatched for better clarity. Environmental bistability states are in a box. (b) Change of pairs sites (stress-tolerant species/fertile site, *ρ*_+1,0_, in blue, competitive species/fertile site, *ρ*_+2,0_, in green) along the fraction of seeds globally dispersed (*δ*). The landscapes corresponding to the asymptotic state of the stochastic spatial model are displayed for *δ* = 0.09 (left) and *δ* = 0.9 (right). Common parameters are *S* = 0, *α_e_* = 0.2. Similar trends are obtained along the stress gradient. (c) Clustering of the stress-tolerant species along the fraction of seeds globally dispersed (*δ*). The clustering coefficient is measured by comparing the density of pairs of stress-tolerant species (*ρ*_+1,+1_) with the expectation under random process 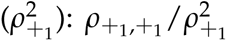. A value above 1 means that species will form clusters more than what is expected by chance. *S* = 0.73 and *α_e_* = 0.2. (d) Change in the thresholds stress of desertification (stress level above which vegetation cover drops to 0) and restoration of vegetation (stress level below which vegetation reappears in desert) along the fraction of seeds globally dispersed (*δ*). *α_e_* = 0.2. The difference between the two lines corresponds to the width of the bistability area.

At higher competition the state at which the community ends up depends on the initial conditions (Fig. 2a hashed gray area). The system is said to be ‘bistable’ as there are two possible system states that can be reached depending on the initial conditions. Those two possible states are: either the two species coexist or the stress-tolerant species ends up alone. The latter result may seem surprising (as we could expect an advantage of the competitive species at high competition) but can be explained as follows. The stress-tolerant species favors the emergence of fertile sites in its neighborhood (via local facilitation) that are weakly accessible to the competitive species because dispersal of seeds by all plants is mainly local (*δ* = 0.1). As a result, starting from an initially low vegetation cover the level of interspecific competition experienced by each species is weak, which favors the dominance of the stress-tolerant species that facilitates the creation of new fertile sites in its neighborhood (see Fig. S1.4). By contrast, when the initial vegetation cover is high, the stress-tolerant species experiences more competition relatively to the competitive one (the latter is less sensitive to competition), which favors the coexistence of the two species. This type of bistability is what we call ‘community bistability’ in what follows. It refers to cases where the different stable states have vegetation, but the species composition of these states differs. The occurrence of this community-scale bistability increases when the two species are both stress-tolerant (*i.e.,* are more sensitive to competition) and have similar strategies (Fig. S1.5a).

As the stress level increases, we observe the following succession of community states: the stress-tolerant species increases in cover until being alone, then the system is bistable between a vegetated and the desert state, before desertification occurs (*i.e.* both species go extinct; see Fig. S1.6 for illustrating transects of Fig. 2a).

We name the bistability between a vegetated and a desert state observed at high-stress levels ‘environmental bistability’. Such bistability has classically been observed in previous models (*e.g.,* Rietkerk and van de Koppel, 1997; von Hardenberg et al., 2001; Kéfi et al., 2007). Environmental bistability emerges from a positive feedback loop between the local vegetation cover and the availability of fertile sites. Specifically, the creation of fertile sites by the stress-tolerant species allows the local regeneration rate of degraded sites (*r* + *f*_0_*q*_+1_ *_|−_* in eq. 5) to outbalance the degradation rate of fertile sites (*d*), which increases recruitment opportunities of vegetation locally (Fig. S1.7). Therefore, when starting from an initially high vegetation cover, the transition is favored toward the creation of new fertile sites, which drives the ecosystem to a vegetated state. By contrast, when stating from an initially low vegetation cover, it drives the ecosystem to a desert state because the degradation rate of fertile sites exceeds their rate of regeneration. Interestingly, the fact that our model contains multiple species allows seeing that the vegetated state of that bistability area can correspond to either the coexistence of the two species (brown hashed area) or to the survival of the stress-tolerant alone (green hashed area on Fig. 2a).

When the dispersal of both species occurs throughout the whole landscape rather than locally (thereby generating a mismatch of scales between dispersal being global and facilitation being local), the competitive species can access more easily fertile sites via the dispersal of its seeds. Consequently, the probability of a competitive species and a fertile site to be neighbors (*ρ*_0,+1_ ) increases (Fig. 2b), while, conversely, the globally dispersed seeds from the stress-tolerant species become less successful at reaching the locally produced fertile sites, and ultimately ends up less spatially clustered (Fig. 2c). Because of these mechanisms, the competitive species ends up out-competing the stress-tolerant species at low stress levels and high interspecific competition (green area in Fig. 2a and Fig. S1.8 for illustration). In addition, this leads to a larger environmental bistability area in case of the mismatch of scales between facilitation and dispersal than when dispersal is local (hashed green area in Fig. 2a; distance between the two curves in Fig. 2d).

So far, the results referred to communities of two species. Next, we increased the species richness of the community.

### Community turnovers in species-richer communities

In multi-species communities, we observe that while the total vegetation cover decreases gradually along the stress gradient (different lines in Fig. 3a), they are important species turnovers, where one or a few species replace the resident community with abrupt changes in the cover of individual species (Fig. 4). At low stress levels, most competitive species dominate communities in term of individual species cover, while as stress level increases resident species are replaced by more stress-tolerant species (as exemplified by the changes in mean community trait, *ψ̄*, in Fig. 3a).

**Figure 3:**
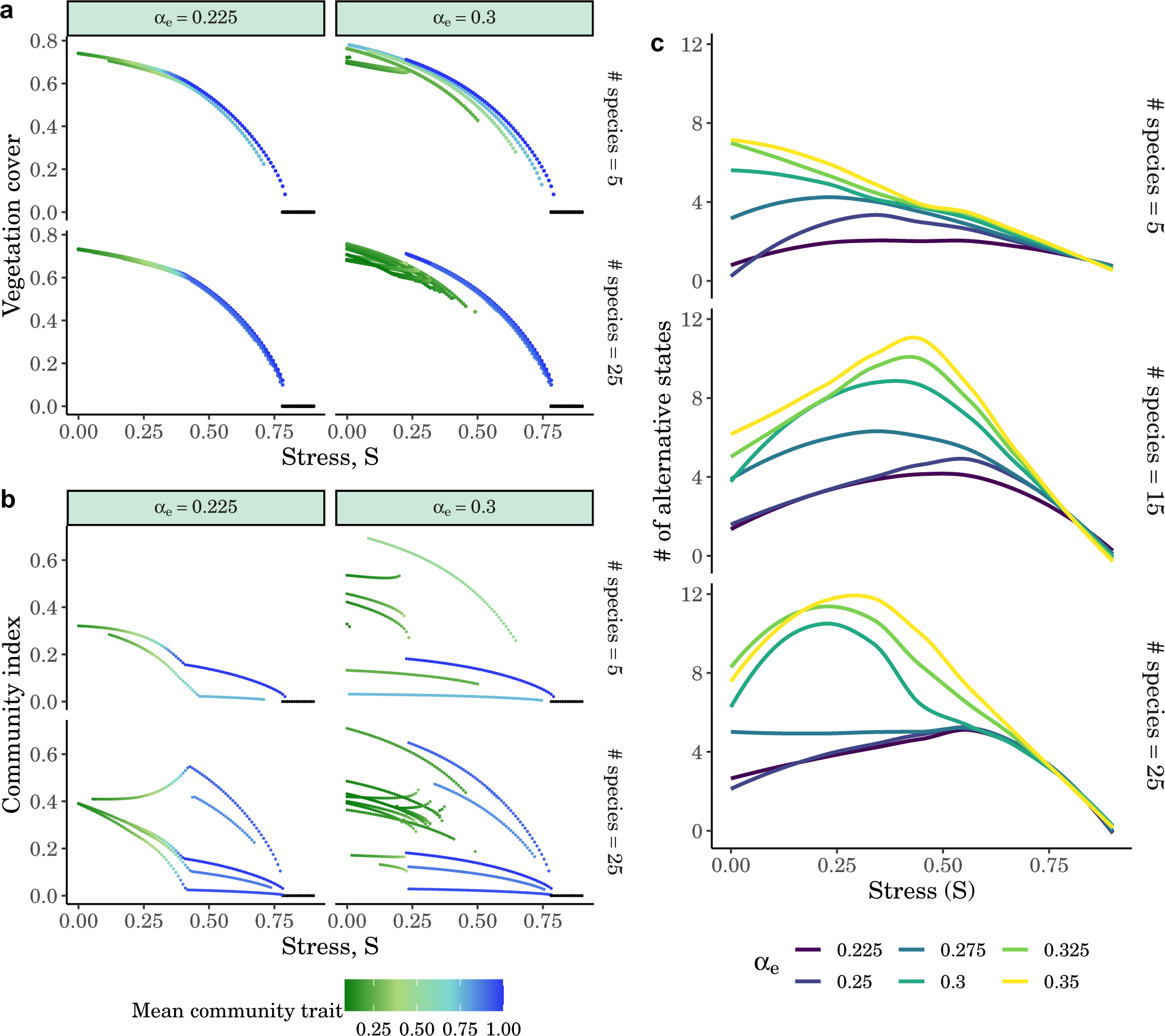
Competition and stress level determine the occurrence of multistability and the number of alternative states. (a) Total vegetation cover for different initial species cover along stress gradient for two levels of species richness (rows, *N* = 5, 25) and two strengths of interspecific interaction (columns, *α_e_* = 0.225, 0.35). The color indicates the community-weighted mean strategy 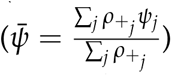. (b) Similar to (a) but with the community index to distinguish communities with similar cover but different species composition. (c) Number of alternative stable states along the stress gradient for varying competition strength and species richness levels. In each panel of community richness (rows), and for each of the interspecific competition strength (*α_e_ ∈ {*0.225, 0.25, 0.275, 0.3, 0.325, 0.35*}* (colors), we counted the number of alternative stable states for each stress level using the number of different species composition at the final state. Lines were smoothed using the *loess* method.

**Figure 4:**
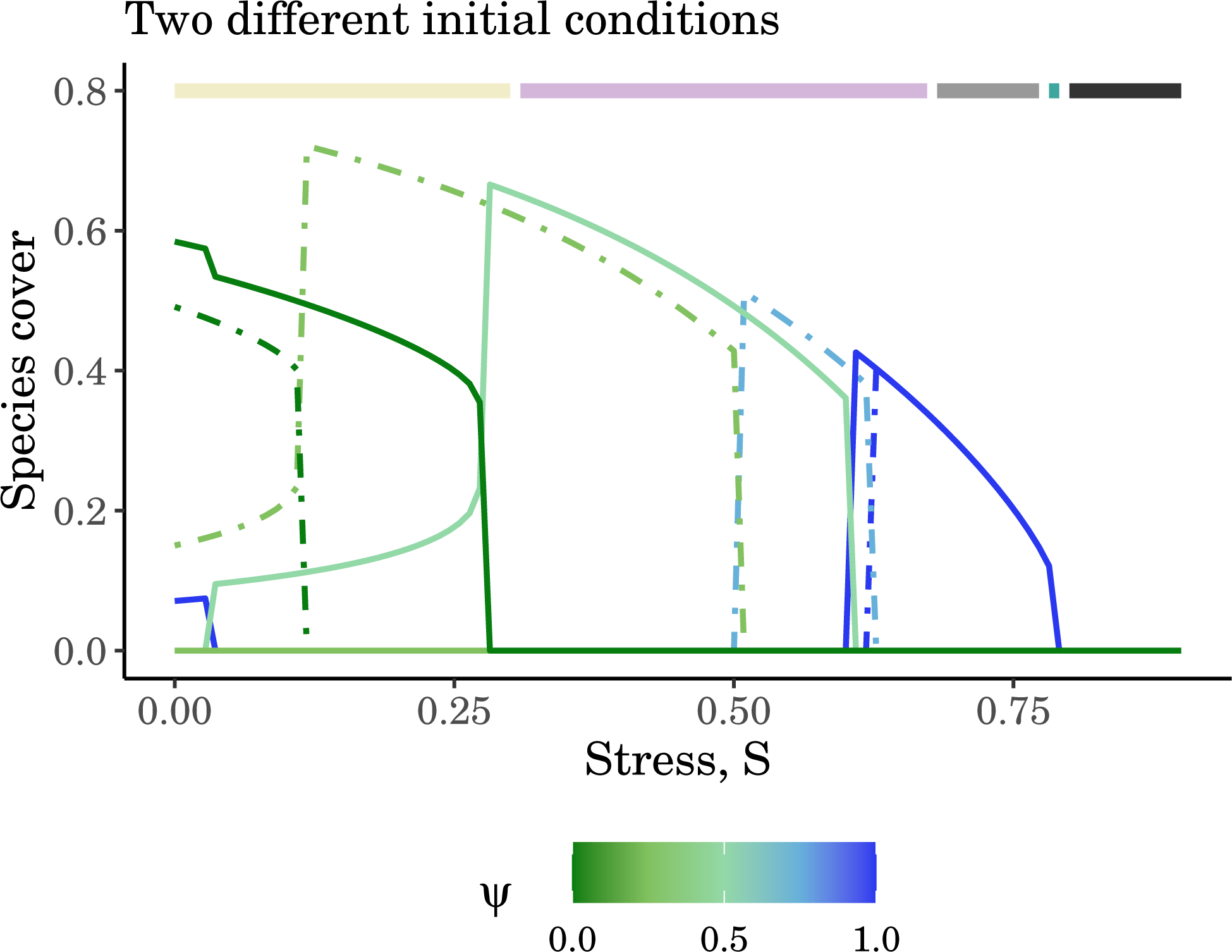
Example dynamics along the stress gradient in species-rich communities. For two different initial conditions (dashed and full lines), we show an example where two different cliques emerge at low-stress conditions (*S <* 0.3) before species become mutually excluding each other as stress level increases (*S ∈ {*0.3, 0.75*}*)). The colored lines above correspond to the different types of multistability (see Fig. 5). Here 5 species are considered in the community. See Fig. S1.11 for other examples.

When varying the initial species cover, community assembly can result in alternative community states with contrasting species compositions as shown by the parallel lines in Fig. 3b. The total number of alternative community states peaks at intermediate levels of stress when the different strategies can invade the landscape (Fig. 3c): at most, we observed about twelve alternative stable states for communities with 15 to 25 species.

### Emergence of different types of community multistability

Depending on the level of stress, the number of coexisting species decreases to one or a few species with increasing competition and stress-level (Figs. 5a, S1.9). In addition to the environmental bistability observed under high stress-levels, we found two types of community multistability.

**Figure 5:**
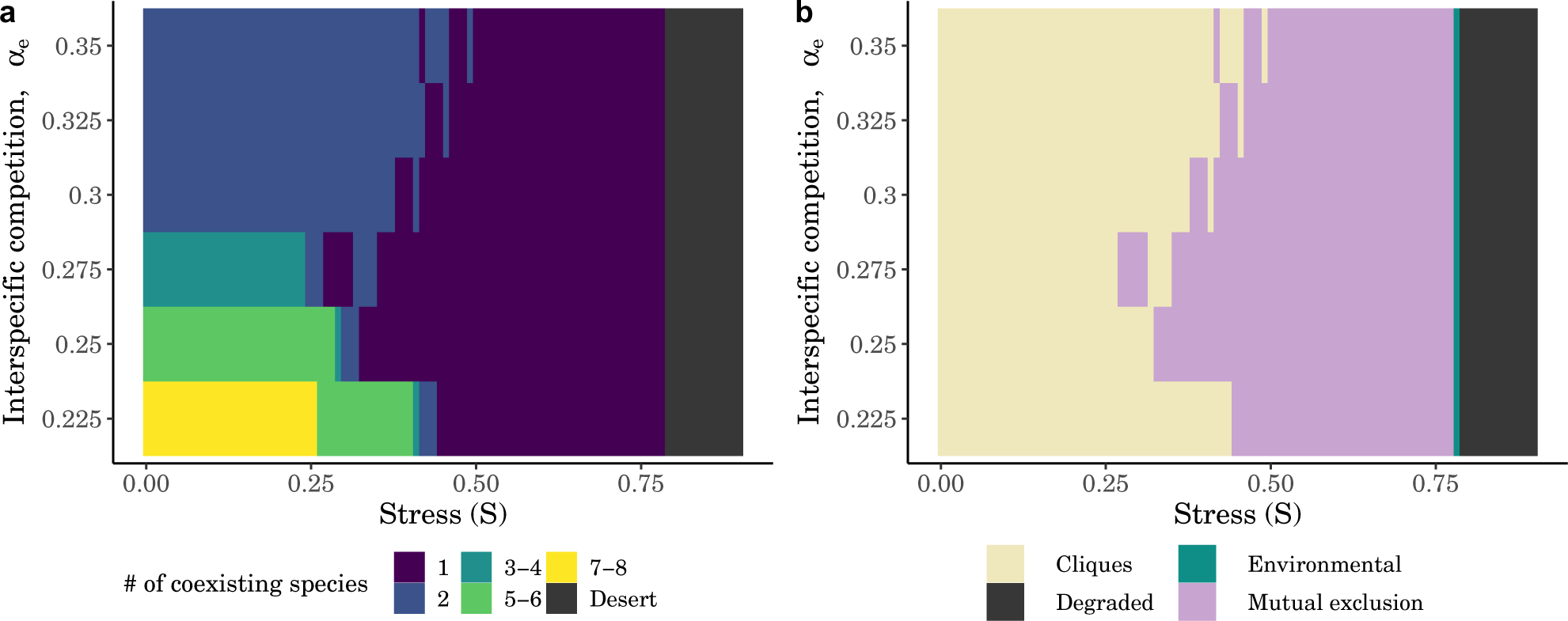
Two types of community multistability emerge depending on the level of stress. (a) Mean number of species observed across all different initial conditions for different levels of interspecific competition (y-axis) and abiotic stress (x-axis). (b) Emergence of two types of community multistability: cliques corresponding to subgroup of coexisting species, and on the other side mutual exclusion states, where species are mutually excluding each other. There is also environmental bistability under higher stress conditions as in the two-species model. See also Fig. S4.5 for a similar figure using the mean-field approach.

Under low-stress conditions (*S <* 0.3), community assembly ends up in sub-groups of coexisting species with different strategies that can co-occur for similar stress conditions (hereafter called ‘cliques’, *sensu* Fried et al., 2016; yellow area in Fig. 5b). The size of these cliques decreases with the level of competition until cliques contain a minimum of two species (Fig. 6a). The composition of these assemblages is not random: they typically host a facilitating species with one or multiple species with more competitive strategies (Fig. 6b, illustrated in Fig. 4). Interestingly, these cliques are maintained by space. Because facilitating plants foster the transition of the degraded sites into fertile ones in their neighborhood, they create favorable conditions for recruitment and thereby buffer against any competitive exclusion from more competitive species (see Appendix S4 for the non-spatial model).

**Figure 6:**
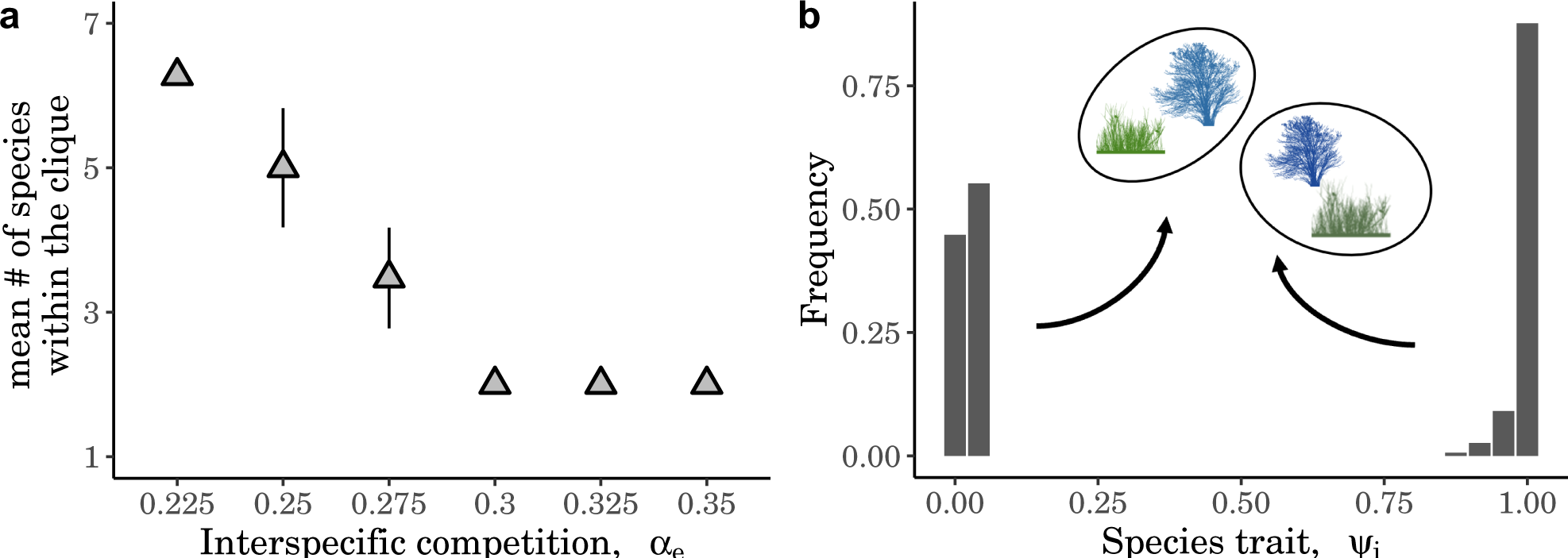
Emerging cliques decrease in size with competition, and are composed of species with constrasting strategies. (a) Number of species coexisting within these cliques along the gradient of competition strength average across all simulations resulting in cliques. (b) Example of distribution of frequency of observation of each strategy across all observed cliques at *S* = 0.1, *α_e_* = 0.35.

In contrast, when abiotic stress increases (*S >* 0.3), we no longer observe cliques; these sub-group of species split, and we observe mutual exclusion states, where only one species with a stress-tolerance strategy dominates the landscape (purple area in Fig. 5b). At extreme levels of stress, only the most stress-tolerant species dominates the landscape while at more intermediate levels of stress, priority effects driven by initial vegetation cover lead to different species with stress-tolerant strategies able to occupy the landscape (Fig. 6b).

Last, we found the two types of community multistability (namely cliques and mutual exclusion states) to have different imprint on the total vegetation cover: because species compete within the cliques, vegetation is on average lower for cliques compared to when there is a single dominant species (Fig. S1.10).

## Discussion

This work investigated how bistability in simple ecological communities expands to multistability in complex communities. In our multi-species plant community model, we found three emergent types of multistability: an environmental bistability between the desert and a vegetated state, as well as two types of community multistability, with either the community composed of small subsets of coexisting species (cliques) or a single dominant species (mutual exclusion) depending on the level of stress. These different states and their possible coexistence can be explained by the different positive feedbacks at play.

### Multistability emerge from different positive feedbacks

Environmental bistability emerges from a positive feedback related to the local facilitation mechanism. By increasing the regeneration of degraded sites, facilitation creates fertile sites in the neighborhood of the most stress-tolerant plants, which can then be recolonized by new plants when reached by seeds. This well described positive feedback fosters vegetation clustering and buffers the effect of stress on vegetation by allowing vegetation to recruit under high stress conditions (Kéfi et al., 2007; Filazzola and Lortie, 2014). When a threshold of stress is reached, this local facilitation is no longer enough to maintain vegetation and desertification occurs with an abrupt loss of vegetation cover (Rietkerk and van de Koppel, 1997).

The two types of community multistability we found also emerge from positive feedbacks but due to species competition. Specifically, the better competitors prevent the recruitment of the weaker ones on the landscape, thereby leaving more space for the recruitment of their own seedlings and contributing to the emergence of positive feedbacks on their recruitment (see Liautaud et al., 2019 for a non-spatial context). When competition is strong, such positive feedback loops can generate strong priority effects, and therefore a dependency to the initial conditions (historical contingency; Fukami, 2015): when a plant species has initially a low cover, the species with higher vegetation cover can inhibit its ability to recruit on fertile sites and to colonize the landscape. By contrast, the same species may persist when starting from an initially high cover (*i.e.,* when it experiences less competition from other species). Hence, each plant species experiences an Allee-like effect, which emerges when interspecific competition is high and leads to alternative community states (Case, 1990; van Nes and Scheffer, 2004).

### Stress level determines the type of community bistability

At low stress levels, we found that the community was organized in small subsets of coexisting species called cliques. In a non-spatial context, cliques have been shown to emerge from the heterogeneity in interaction strengths (Bunin, 2017; Aguadé-Gorgorió et al., 2023). Such heterogeneity prevents any competitive hierarchy of species by generating instansivity (Kerr et al., 2002; Allesina and Levine, 2011). By contrast, in our model, the emergence of cliques was promoted by space, and not by variation in species interaction strengths. Indeed, because seeds are mainly dispersed locally in drylands, facilitation mostly benefits facilitating species, which have more suitable sites in their direct neighborhood on which seeds can recruit (*i.e.,* positive niche construction; Odling-Smee et al., 1996). This mechanism buffers against competitive exclusion of the most facilitating species in the community. Specifically, at equilibrium, local positive niche construction effects out-balance the negative effects from interspecific competition on the facilitating species and promote the emergence of small groups of coexisting species with at least one stress-tolerant and one competitive species. However, when dispersal is global or when there is no spatial structure, facilitation can be seen as a cooperative trait that benefits all species, and our results indicate that it hinders the persistence of the facilitating species and promotes mutual exclusion states, where only one species dominates the landscape (Appendix S4). In such a scenario, facilitation may therefore not be selected over evolutionary timescales (Kéfi et al., 2008).

At high levels of stress, cliques break into mutual exclusion states, each characterized by the dominance of a single facilitating species. This means that, under highly stressed environments, only one species with a high stress-tolerance dominates the landscape in our model. This result echoes a recent theory coined by Zobel et al. (2023) to explain the patterns of coexistence in communities: when positive niche construction dominates the ecosystem, it can lead to the dominance of a single species. Indeed, in any stressed ecosystem (with high aridity, salt levels, or low pH), high stress levels limit the recruitment of new individuals, leaving space for species that actively modify their environment and buffer the local stress level. In high latitude peat bogs, for instance, low pH and moisture favor the dominance of a moss species (*Sphagnum*; Clymo, 1984). This moss species further shapes soil conditions by increasing acidity, limiting decomposition and the growth of other species, together promoting its ecosystem-level dominance. In drylands, such a decrease in species and functional diversity can be expected as aridity increases. Facilitation by nurse species allows species to persist beyond their niche until aridity reaches a threshold, where facilitation cannot sustain the less tolerant species anymore, and species diversity decreases rapidly. Such a threshold has been estimated to happen at an aridity index of about 0.8 (Berdugo et al., 2020). In our simulation results, only one single stress-tolerant species is able to persist at high stress levels but our model only considers one trait axis. Taking multiple traits into account would likely lead to the coexistence of a small group of species dominated by a stress-tolerant species rather than a single species, which in better agreement with projection of drylands in a warmer climate (Frenette Dussault et al., 2013).

### Community turnover in drylands

We found that as the stress level increases, there are abrupt shifts in community composition. Such abrupt changes in individual species cover are due to strong interspecific competition and the emergence of alternative states in our model. Specifically, species are highly integrated due to the positive feedbacks emerging from competition that clear boundaries separating each species emerge along the stress-gradient (Liautaud et al., 2019). Furthermore, more stress-tolerant strategies rapidly replace more competitive ones as aridity increases, leading to important community turnovers along the stress gradient before desertification. Current knowledge on drylands supports these findings. For instance, because of desertification, the functional composition of communities is drastically changing: stress-tolerant species (*e.g.,* shrubs) becomes more frequently observed (Frenette Dussault et al., 2013; Berdugo et al., 2020), while the cover of perennial herbs or grasses, that are overall less tolerant to stress, tend to decrease (Munson et al., 2012). With the current increase in climate variability worldwide and notably in drylands (Mirzabaev et al., 2022), we expect these community turnovers to increase in frequency as they have been shown to be promoted by the variability of temperature and precipitations (Ulrich et al., 2014). Given the strong competition between species for water, the succession of plants along the stress gradient may therefore be characterized by rapid species turnover.

So far, current evaluation of dryland degradation is highly based on assessing changes in overall vegetation cover. The use of remote sensing provide important tools to follow drylands in time and space (Kéfi et al., 2014; Majumder et al., 2019). However, this approach is blind to changes in species composition. Because changes in functional composition of plant communities might buffer the stability of drylands (*e.g.,* by increasing stress-tolerant species), getting information about functional composition of plant communities might provide better indicators of ecosystem state change (Gross et al., 2013). In particular, identifying thresholds of aridity associated with functional reorganization of dryland plant communities will complement current thresholds associated with vegetation cover decline and loss of soil functioning (Berdugo et al., 2020).

### Perspectives

While our model allows to derive some prediction and mechanisms for plant communities in a warmer world, we ignored the role of grazing, yet considered to be one of the main drivers of desertification (Asner et al., 2004). Grazing has been shown to decrease species diversity (Merdas et al., 2021) and shift community composition to plant strategies investing more energy in below-ground structures (Nathan et al., 2016). Together, given that grazing and aridity can interact (Gao et al., 2018), that plants exhibit plastic responses to stress by switching from tolerance to avoidance when reaching their physiological limits (Kramp et al., 2022), we expect community succession in drylands to be more complex than predicted by our model. This calls to future studies integrating abiotic and biotic pressures to better understand the succession of plant strategies in drier drylands.

By scaling-up our understanding of bistability in stressed ecosystems from one species to the community level, our study sets bases to better understand how different types of multistability emerge in species and interaction-rich communities. Pursuing to quantify the dynamical and emergent properties resulting from community-feedbacks will help to widen our understanding of how the multiplexity of feedback loops determines our understanding of the stability of ecological systems and more generally, of complex systems with many interacting entities, such as gene regulatory networks, or gut microbiome interactions.

## Acknowledgments

B.P acknowledges a doctoral fellowship from the chaire Modélisation Mathématique et Biodiversité of VEOLIA-Ecole Polytechnique-MNHN.

## Author contribution

All authors designed the study. B.P performed research and wrote the first draft. All authors substantially revised the manuscript.

## Literature Cited

Aguadé-Gorgorió, G., J.-F. Arnoldi, M. Barbier, and S. Kéfi. 2023. A taxonomy of multiple stable states in complex ecological communities. bioRxiv, page 2023.08.30.555051. URL http://biorxiv.org/content/early/2023/08/31/2023.08.30.555051.abstract.

Aguiar, M. R., O. E. Sala, and M. R. Aguiar. 1994. Competition, facilitation, seed distribution and the origin of patches in a Patagonian steppe. Oikos, 70:26. URL https://www.jstor.org/stable/3545695?origin=crossref.

Allesina, S. and J. M. Levine. 2011. A competitive network theory of species diversity. Proceedings of the National Academy of Sciences, 108:5638–5642. URL https://pnas.org/doi/full/10.1073/pnas.1014428108.

Angert, A., T. Huxman, P. Chesson, and D. Venable. 2009. Functional tradeoffs determine species coexistence via the storage effect. Proceedings of the National Academy of Sciences of the United States of America, 106:11641–5.

Armas, C. and F. I. Pugnaire. 2005. Plant interactions govern population dynamics in a semiarid plant community. Journal of Ecology, 93:978–989. URL https://onlinelibrary.wiley.com/doi/10.1111/j.1365-2745.2005.01033.x.

Asner, G. P., A. J. Elmore, L. P. Olander, R. E. Martin, and A. T. Harris. 2004. Grazing systems, ecosystem responses, and global change. Annual Review of Environment and Resources, 29:261–299. URL https://www.annualreviews.org/doi/10.1146/annurev.energy.29.062403.102142.

Aubier, T. G. 2020. Positive density dependence acting on mortality can help maintain species-rich communities. eLife, 9:e57788. URL https://elifesciences.org/articles/57788.

Berdugo, M., M. Delgado-Baquerizo, S. Soliveres, R. Hernández-Clemente, Y. Zhao, J. J. Gaitán, N. Gross, H. Saiz, V. Maire, A. Lehmann, M. C. Rillig, R. V. Solé, and F. T. Maestre. 2020. Global ecosystem thresholds driven by aridity. Science, 367:787–790. URL https://www.science.org/doi/10.1126/science.aay5958.

Bunin, G. 2017. Ecological communities with Lotka-Volterra dynamics. Physical Review E, 95:042414. URL http://link.aps.org/doi/10.1103/PhysRevE.95.042414.

Burns, J. H. and S. Y. Strauss. 2011. More closely related species are more ecologically similar in an experimental test. Proceedings of the National Academy of Sciences, 108:5302–5307. URL https://pnas.org/doi/full/10.1073/pnas.1013003108.

Burrell, A. L., J. P. Evans, and M. G. De Kauwe. 2020. Anthropogenic climate change has driven over 5 million km2 of drylands towards desertification. Nature Communications, 11:3853. URL http://www.nature.com/articles/s41467-020-17710-7.

Butterfield, B. J. and J. M. Briggs. 2011. Regeneration niche differentiates functional strategies of desert woody plant species. Oecologia, 165:477–487. URL http://link.springer.com/10.1007/s00442-010-1741-y.

Cahill, J. F., S. W. Kembel, E. G. Lamb, and P. A. Keddy. 2008. Does phylogenetic relatedness influence the strength of competition among vascular plants? Perspectives in Plant Ecology, Evolution and Systematics, 10:41–50. URL https://linkinghub.elsevier.com/retrieve/pii/S1433831907000479.

Callaway, R. 2007. Positive interactions and interdependence in plant communities. Springer Netherlands.

Case, T. J. 1990. Invasion resistance arises in strongly interacting species-rich model competition communities. Proceedings of the National Academy of Sciences, 87:9610– 9614. URL https://pnas.org/doi/full/10.1073/pnas.87.24.9610.

Centeno, M. A., P. W. Callahan, P. A. Larcey, and T. S. Patterson. 2023. How worlds collapse: What history, systems, and complexity can teach us about our modern world and fragile future. Taylor & Francis.

Choler, P., R. Michalet, Ragan, , and C. MR. 2001. Facilitation and competition on gradients in alpine plant communities. Ecology, 82:3295–3308.

Clymo, R. S. 1984. Sphagnum-dominated peat bog: a naturally acid ecosystem. Philosophical Transactions of the Royal Society of London. B, Biological Sciences, 305:487–499. URL https://royalsocietypublishing.org/doi/10.1098/rstb.1984.0072.

Danet, A., F. D. Schneider, F. Anthelme, and S. Kéfi. 2021. Indirect facilitation drives species composition and stability in drylands. Theoretical Ecology, 14:189–203. URL https://link.springer.com/10.1007/s12080-020-00489-0.

De Roos, A. M. and L. Persson. 2002. Size-dependent life-history traits promote catastrophic collapses of top predators. Proceedings of the National Academy of Sciences, 99:12907– 12912. URL https://pnas.org/doi/full/10.1073/pnas.192174199.

DeAngelis, D. L., W. M. Post, and C. C. Travis. 1986. Positive feedback in natural systems. Number volume 15 in Biomathematics, Springer-Verlag, Berlin Heidelberg New York Tokyo.

Downing, A. S., E. H. van Nes, W. M. Mooij, and M. Scheffer. 2012. The resilience and resistance of an ecosystem to a collapse of diversity. PLoS ONE, 7:e46135. URL https://dx.plos.org/10.1371/journal.pone.0046135.

Díaz, S., J. Kattge, J. H. C. Cornelissen, I. J. Wright, S. Lavorel, S. Dray, B. Reu, M. Kleyer, C. Wirth, I. Colin Prentice, E. Garnier, G. Bönisch, M. Westoby, H. Poorter, P. B. Reich, A. T. Moles, J. Dickie, A. N. Gillison, A. E. Zanne, J. Chave, S. Joseph Wright, S. N. Sheremet’ev, H. Jactel, C. Baraloto, B. Cerabolini, S. Pierce, B. Shipley, D. Kirkup, F. Casanoves, J. S. Joswig, A. Günther, V. Falczuk, N. Rüger, M. D. Mahecha, and L. D. Gorné. 2016. The global spectrum of plant form and function. Nature, 529:167–171. URL http://www.nature.com/articles/nature16489.

Fagundes, M. V., R. S. Oliveira, C. R. Fonseca, and G. Ganade. 2022. Nurse-target functional match explains plant facilitation strength. Flora, 292:152061. URL https://linkinghub.elsevier.com/retrieve/pii/S0367253022000585.

Filazzola, A. and C. J. Lortie. 2014. A systematic review and conceptual framework for the mechanistic pathways of nurse plants. Global Ecology and Biogeography, 23:1335–1345. URL https://onlinelibrary.wiley.com/doi/10.1111/geb.12202.

Fllner, S. and A. Shmida. 1981. Why are adaptations for long-range seed dispersal rare in desert plants? Oecologia, 51:133–144. URL http://link.springer.com/10.1007/BF00344663.

Flores, J. and E. Jurado. 2003. Are nurse-protégé interactions more common among plants from arid environments? Journal of Vegetation Science, 14:911–916. URL https://onlinelibrary.wiley.com/doi/10.1111/j.1654-1103.2003.tb02225.x.

Fontaine, C., P. R. Guimarães, S. Kéfi, N. Loeuille, J. Memmott, W. H. van der Putten, F. J. F. van Veen, and E. Thébault. 2011. The ecological and evolutionary implications of merging different types of networks: Merging networks with different interaction types. Ecology Letters, 14:1170–1181. URL https://onlinelibrary.wiley.com/doi/10.1111/j.14610248.2011.01688.x.

Frenette Dussault, C., B. Shipley, D. Meziane, and Y. Hingrat. 2013. Trait-based climate change predictions of plant community structure in arid steppes. Journal of Ecology, 101:484–492.

Fried, Y., D. A. Kessler, and N. M. Shnerb. 2016. Communities as cliques. Scientific Reports, 6:35648. URL http://www.nature.com/articles/srep35648.

Fukami, T. 2015. Historical contingency in community assembly: Integrating niches, species pools, and priority Effects. Annual Review of Ecology, Evolution, and Systematics, 46:1–23. URL https://www.annualreviews.org/doi/10.1146/annurev-ecolsys-110411-160340.

Gao, S., Z. Zheng, Y. Wang, L. Liu, N. Zhao, and Y. Gao. 2018. Drought and grazing drive the retrogressive succession by changing the plant-plant interaction of the main species in Inner Mongolia Steppe. Ecology and Evolution, 8:11954–11963. URL https://onlinelibrary.wiley.com/doi/10.1002/ece3.4652.

Gilad, E., M. Shachak, and E. Meron. 2007. Dynamics and spatial organization of plant communities in water-limited systems. Theoretical Population Biology, 72:214–230. URL https://linkinghub.elsevier.com/retrieve/pii/S0040580907000603.

Gilpin, M. E. and T. J. Case. 1976. Multiple domains of attraction in competition communities. Nature, 261:40–42. URL 10.1038/261040a0.

Graff, P. and M. R. Aguiar. 2017. Do species’ strategies and type of stress predict net positive effects in an arid ecosystem? Ecology, 98:794–806. URL https://onlinelibrary.wiley.com/doi/10.1002/ecy.1703.

Gross, K. 2008. Positive interactions among competitors can produce species-rich communities. Ecology Letters, 11:929–936. URL https://onlinelibrary.wiley.com/doi/10.1111/j.1461-0248.2008.01204.x.

Gross, N., L. Börger, S. I. Soriano-Morales, Y. Le Bagousse-Pinguet, J. L. Quero, M. García-Gómez, E. Valencia-Gómez, and F. T. Maestre. 2013. Uncovering multiscale effects of aridity and biotic interactions on the functional structure of Mediterranean shrublands. Journal of Ecology, 101:637–649. URL https://onlinelibrary.wiley.com/doi/ 10.1111/1365-2745.12063.

He, Q., B. Cui, M. D. Bertness, and Y. An. 2012. Testing the importance of plant strategies on facilitation using congeners in a coastal community. Ecology, 93:2023–2029. URL http://www.jstor.org/stable/41739260.

Janssen, M., T. Kohler, and M. Scheffer. 2003. Sunk-cost effects and vulnerability to collapse in ancient societies. Curent, 44:722–728.

Karatayev, V. A., M. L. Baskett, and E. H. van Nes. 2023. The potential for alternative stable states in food webs depends on feedback mechanism and trait diversity. The American Naturalist, 202:260–275. URL 10.1086/725421. Publisher: The University of Chicago Press.

Kerr, B., M. A. Riley, M. W. Feldman, and B. J. M. Bohannan. 2002. Local dispersal promotes biodiversity in a real-life game of rock–paper–scissors. Nature, 418:171–174. URL https://www.nature.com/articles/nature00823.

Kramp, R. E., P. Liancourt, M. M. Herberich, L. Saul, S. Weides, K. Tielbörger, and M. Májeková. 2022. Functional traits and their plasticity shift from tolerant to avoidant under extreme drought. Ecology. URL https://onlinelibrary.wiley.com/doi/10.1002/ecy.3826.

Kéfi, S., M. Baalen, M. Rietkerk, and M. Loreau. 2008. Evolution of local facilitation in arid ecosystems. The American Naturalist, 172:E1–E17. URL https://www.journals.uchicago.edu/doi/10.1086/588066.

Kéfi, S., E. L. Berlow, E. A. Wieters, S. A. Navarrete, O. L. Petchey, S. A. Wood, A. Boit, L. N. Joppa, K. D. Lafferty, R. J. Williams, N. D. Martinez, B. A. Menge, C. A. Blanchette, A. C. Iles, and U. Brose. 2012. More than a meal. . . integrating non-feeding interactions into food webs. Ecology Letters, 15:291–300. URL https://onlinelibrary.wiley.com/doi/ 10.1111/j.1461-0248.2011.01732.x.

Kéfi, S., V. Guttal, W. A. Brock, S. R. Carpenter, A. M. Ellison, V. N. Livina, D. A. Seekell, M. Scheffer, E. H. van Nes, and V. Dakos. 2014. Early warning signals of ecological transitions: methods for spatial patterns. PLOS ONE, 9:e92097. URL 10.1371/journal.pone.0092097. Publisher: Public Library of Science.

Kéfi, S., M. Holmgren, and M. Scheffer. 2016a. When can positive interactions cause alternative stable states in ecosystems? Functional Ecology, 30:88–97. URL https://onlinelibrary.wiley.com/doi/10.1111/1365-2435.12601.

Kéfi, S., V. Miele, E. A. Wieters, S. A. Navarrete, and E. L. Berlow. 2016b. How structured is the entangled bank? The surprisingly simple organization of multiplex ecological networks leads to increased persistence and resilience. PLOS Biology, 14:e1002527. URL https://dx.plos.org/10.1371/journal.pbio.1002527.

Kéfi, S., M. Rietkerk, M. van Baalen, and M. Loreau. 2007. Local facilitation, bistability and transitions in arid ecosystems. Theoretical Population Biology, 71:367–379. URL https://linkinghub.elsevier.com/retrieve/pii/S0040580906001250.

Kéfi, S., C. Saade, E. L. Berlow, J. S. Cabral, and E. A. Fronhofer. 2022. Scaling up our understanding of tipping points. Philosophical Transactions of the Royal Society B: Biological Sciences, 377:20210386. URL https://royalsocietypublishing.org/doi/10.1098/rstb.2021.0386.

Lenton, T. M., H. Held, E. Kriegler, J. W. Hall, W. Lucht, S. Rahmstorf, and H. J. Schellnhuber. 2008. Tipping elements in the Earth’s climate system. Proceedings of the National Academy of Sciences, 105:1786–1793. URL http://www.pnas.org/cgi/doi/10.1073/pnas.0705414105.

Lever, J. J., I. A. de Leemput, E. Weinans, R. Quax, V. Dakos, E. H. Nes, J. Bascompte, and M. Scheffer. 2020. Foreseeing the future of mutualistic communities beyond collapse. Ecology Letters, 23:2–15. URL https://onlinelibrary.wiley.com/doi/10.1111/ele.13401.

Lever, J. J., E. H. van Nes, M. Scheffer, and J. Bascompte. 2014. The sudden collapse of pollinator communities. Ecology Letters, 17:350–359. URL https://onlinelibrary.wiley.com/doi/10.1111/ele.12236.

Liancourt, P., R. M. Callaway, and R. Michalet. 2005. Stress tolerance and competitive-response ability determine the outcome of biotic interactions. Ecology, 86:1611–1618. URL http://www.jstor.org/stable/3450786.

Liautaud, K., E. H. van Nes, M. Barbier, M. Scheffer, and M. Loreau. 2019. Super-organisms or loose collections of species? A unifying theory of community patterns along environmental gradients. Ecology Letters, page ele.13289. URL https://onlinelibrary.wiley.com/doi/10.1111/ele.13289.

MacArthur, R. and R. Levins. 1967. The limiting similarity, convergence, and divergence of coexisting species. The American Naturalist, 101:377–385. URL https://www.journals.uchicago.edu/doi/10.1086/282505.

Maestre, F. T., J. L. Quero, N. J. Gotelli, A. Escudero, V. Ochoa, M. Delgado-Baquerizo, M. García-Gómez, M. A. Bowker, S. Soliveres, C. Escolar, P. García-Palacios, M. Berdugo, E. Valencia, B. Gozalo, A. Gallardo, L. Aguilera, T. Arredondo, J. Blones, B. Boeken, D. Bran, A. A. Conceição, O. Cabrera, M. Chaieb, M. Derak, D. J. Eldridge, C. I. Espinosa, A. Florentino, J. Gaitán, M. G. Gatica, W. Ghiloufi, S. Gómez-González, J. R. Gutiérrez, R. M. Hernández, X. Huang, E. Huber-Sannwald, M. Jankju, M. Miriti, J. Monerris, R. L. Mau, E. Morici, K. Naseri, A. Ospina, V. Polo, A. Prina, E. Pucheta, D. A. Ramírez-Collantes, R. Romão, M. Tighe, C. Torres-Díaz, J. Val, J. P. Veiga, D. Wang, and E. Zaady. 2012. Plant species richness and ecosystem multifunctionality in global drylands. Science, 335:214–218. URL https://www.science.org/doi/10.1126/science.1215442.

Majumder, S., K. Tamma, S. Ramaswamy, and V. Guttal. 2019. Inferring critical thresholds of ecosystem transitions from spatial data. Ecology, 100. URL https://onlinelibrary.wiley.com/doi/10.1002/ecy.2722.

Merdas, S., Y. Kouba, T. Mostephaoui, Y. Farhi, and H. Chenchouni. 2021. Livestock grazing-induced large-scale biotic homogenization in arid Mediterranean steppe rangelands. Land Degradation & Development, 32:5099–5107. URL https://onlinelibrary.wiley.com/doi/10.1002/ldr.4095.

Mirzabaev, A., L. Stringer, T. Benjaminsen, P. Gonzalez, R. Harris, M. Jafari, N. Stevens, C. Tirado, and S. Zakieldeen. 2022. Cross-Chapter Paper 3 Deserts, Semiarid Areas and Desertification. In H. O. Pörtner, D. C. Roberts, M. Tignor, E. S. Poloczanska, K. Minten-beck, A. Alegría, M. Craig, S. Langsdorf, S. Löschke, V. Möller, A. Okem, and B. Rama, editors, Climate Change 2022 Impacts, Adaptation and Vulnerability. Contribution of Working Group II to the Sixth Assessment Report of the Intergovernmental Panel on Climate Change, pages 2195–2231. Cambridge University Press, Cambridge, UK and New York, USA. Type: Book Section.

Mougi, A. and M. Kondoh. 2012. Diversity of interaction types and ecological community stability. Science, 337:349–351. URL https://www.sciencemag.org/lookup/doi/10.1126/science.1220529.

Munson, S. M., R. H. Webb, J. Belnap, J. Andrew Hubbard, D. E. Swann, and S. Rutman. 2012. Forecasting climate change impacts to plant community composition in the Sonoran Desert region. Global Change Biology, 18:1083–1095. URL https://onlinelibrary.wiley.com/doi/10.1111/j.1365-2486.2011.02598.x.

Nathan, J., Y. Osem, M. Shachak, and E. Meron. 2016. Linking functional diversity to resource availability and disturbance: a mechanistic approach for water-limited plant communities. Journal of Ecology, 104:419–429. URL 10.1111/1365-2745.12525. Publisher: John Wiley & Sons, Ltd.

Nunes, A., M. Köbel, P. Pinho, P. Matos, F. d. Bello, O. Correia, and C. Branquinho. 2017. Which plant traits respond to aridity? A critical step to assess functional diversity in Mediterranean drylands. Agricultural and Forest Meteorology, 239:176–184. URL https://linkinghub.elsevier.com/retrieve/pii/S0168192317301120.

Odling-Smee, F. J., K. N. Laland, and M. W. Feldman. 1996. Niche construction. The American Naturalist, 147:641–648. Publisher: University of Chicago Press.

Odling-Smee, J., D. H. Erwin, E. P. Palkovacs, M. W. Feldman, and K. N. Laland. 2013. Niche construction theory: A practical guide for ecologists. The Quarterly Review of Biology, 88:3–28. URL https://www.journals.uchicago.edu/doi/10.1086/669266.

Pichon, B., S. Kéfi, N. Loeuille, I. Lajaaiti, and I. Gounand. 2023. Integrating ecological feedbacks across scales and levels of organization. EcoEvoRxiv.

Pocock, M. J. O., D. M. Evans, and J. Memmott. 2012. The robustness and restoration of a network of ecological networks. Science, 335:973–977. URL https://www.sciencemag.org/lookup/doi/10.1126/science.1214915.

Pueyo, Y., S. Kefi, C. L. Alados, and M. Rietkerk. 2008. Dispersal strategies and spatial organization of vegetation in arid ecosystems. Oikos, 117:1522–1532. URL https://onlinelibrary.wiley.com/doi/10.1111/j.0030-1299.2008.16735.x.

Rial, J. A., R. A. Pielke Sr., M. Beniston, M. Claussen, J. Canadell, P. Cox, H. Held, N. de Noblet-Ducoudré, R. Prinn, J. F. Reynolds, and J. D. Salas. 2004. Nonlinearities, feedbacks and critical thresholds within the earth’s climate system. Climatic Change, 65:11–38. URL http://link.springer.com/10.1023/B:CLIM.0000037493.89489.3f.

Rietkerk, M. 2004. Self-organized patchiness and catastrophic shifts in ecosystems. Science, 305:1926–1929. URL https://www.sciencemag.org/lookup/doi/10.1126/science.1101867.

Rietkerk, M. and J. van de Koppel. 1997. Alternate stable states and threshold effects in semi-arid grazing systems. Oikos, 79:69. URL https://www.jstor.org/stable/3546091?origin=crossref.

Ruppert, J. C., K. Harmoney, Z. Henkin, H. A. Snyman, M. Sternberg, W. Willms, and A. Linstädter. 2015. Quantifying drylands’ drought resistance and recovery: the importance of drought intensity, dominant life history and grazing regime. Global Change Biology, 21:1258–1270. URL https://onlinelibrary.wiley.com/doi/10.1111/gcb.12777.

Scheffer, M. 2009. Critical Transitions in Nature and Society. Princeton University Press.

Scheffer, M., S. Carpenter, J. A. Foley, C. Folke, and B. Walker. 2001. Catastrophic shifts in ecosystems. Nature, 413:591–596. URL http://www.nature.com/articles/35098000.

Schlesinger, W. H., J. F. Reynolds, G. L. Cunningham, L. F. Huenneke, W. M. Jarrell, R. A. Virginia, and W. G. Whitford. 1990. Biological feedbacks in global desertification. Science, 247:1043–1048. URL https://www.science.org/doi/10.1126/science.247.4946.1043.

Ulrich, W., S. Soliveres, F. T. Maestre, N. J. Gotelli, J. L. Quero, M. Delgado-Baquerizo, M. A. Bowker, D. J. Eldridge, V. Ochoa, B. Gozalo, E. Valencia, M. Berdugo, C. Escolar, M. García-Gómez, A. Escudero, A. Prina, G. Alfonso, T. Arredondo, D. Bran, O. Cabrera, A. P. Cea, M. Chaieb, J. Contreras, M. Derak, C. I. Espinosa, A. Florentino, J. Gaitán, V. G. Muro, W. Ghiloufi, S. Gómez-González, J. R. Gutiérrez, R. M. Hernández, E. Huber-Sannwald, M. Jankju, R. L. Mau, F. M. Hughes, M. Miriti, J. Monerris, M. Muchane, K. Naseri, E. Pucheta, D. A. Ramírez-Collantes, E. Raveh, R. L. Romão, C. Torres-Díaz, J. Val, J. P. Veiga, D. Wang, X. Yuan, and E. Zaady. 2014. Climate and soil attributes determine plant species turnover in global drylands. Journal of Biogeography, 41:2307– 2319. URL https://onlinelibrary.wiley.com/doi/10.1111/jbi.12377.

Valiente-Banuet, A., A. V. Rumebe, M. Verdú, and R. M. Callaway. 2006. Modern Qua-ternary plant lineages promote diversity through facilitation of ancient Tertiary lineages. Proceedings of the National Academy of Sciences, 103:16812–16817. URL https://pnas.org/doi/full/10.1073/pnas.0604933103.

van Nes, E. and M. Scheffer. 2004. Large species shifts triggered by small forces. The American Naturalist, 164:255–266. URL http://www.journals.uchicago.edu/doi/10.1086/422204.

von Hardenberg, J., E. Meron, M. Shachak, and Y. Zarmi. 2001. Diversity of vegetation patterns and desertification. Physical Review Letters, 87:198101. URL https://link.aps.org/doi/10.1103/PhysRevLett.87.198101.

Zhang, R. and K. Tielbörger. 2019. Facilitation from an intraspecific perspective – stress tolerance determines facilitative effect and response in plants. New Phytologist, 221:2203– 2212. URL https://onlinelibrary.wiley.com/doi/10.1111/nph.15528.

Zobel, M., M. Moora, M. Pärtel, M. Semchenko, L. Tedersoo, M. Öpik, and J. Davison. 2023. The multiscale feedback theory of biodiversity. Trends in Ecology & Evolution, 38:171–182. URL https://linkinghub.elsevier.com/retrieve/pii/S0169534722002282.

