## Supplementary material for "The interplay of facilitation and competition drives the emergence of multistability in dryland plant communities": SI_text

**S1 Additional figures referenced in the main text** **S3**

**S2 Relaxing the shape of the trade-off** **S15**

**S3 Supplementary methods** **S19**

**S4 Mean-field model** **S23**

**S5 Pair approximation model** **S30**

### S1 Additional figures referenced in the main text

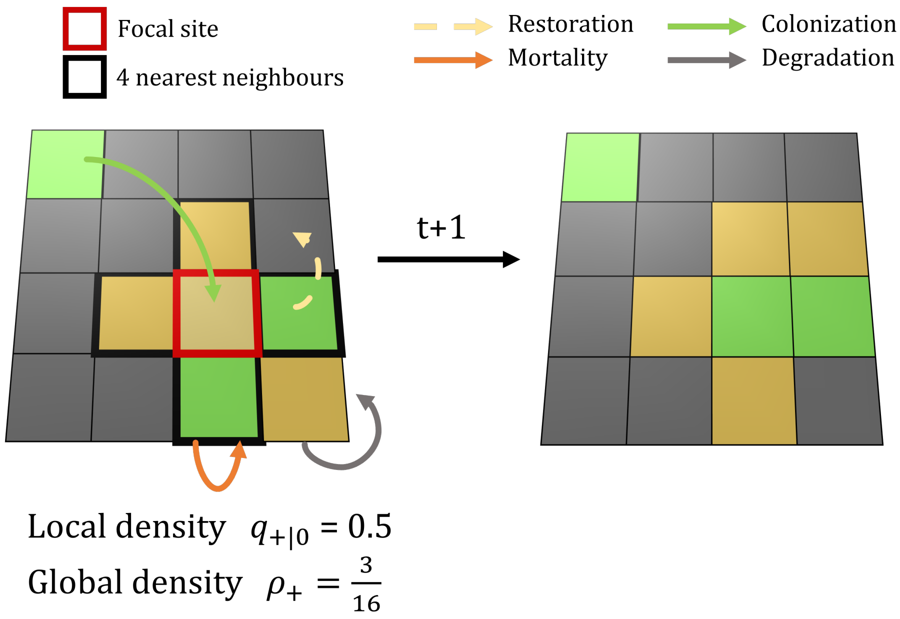


Figure S1.1: **Definitions of the local and global variables and illustration of the 4 possible transitions between the states.**
In a hypothetical $4 \times 4$ landscape, there are two main quantities tracked by the model. First, the global densities of sites (here vegetation $\rho_+$). Here, in this case, as we display only one plant species ($N = 1$), we have $\rho_{+_1} = \sum_{\text{site}+} /16$. In addition, for a given focal site, we can compute the local density of vegetation in the 4 nearest neighbors of the focal site. Here we illustrate that with $q_{+|0}$ with a fertile site as the focal site. We also show the 4 possible transitions in the model: a site of vegetation can colonize and recruit on a fertile site (green arrow) or go extinct and give a fertile site by mortality (orange arrow). Fertile sites are formed by restoration. The rate of restoration is increased by the local density of vegetation in the 4 nearest sites of a degraded site (dashed yellow arrow). Finally, fertile sites can give degraded sites by degradation of the sites (grey arrow). These probabilities of transition between each of the $N + 2$ states do not change with time. Hence, at any time $t$, the probability of a given site changing in another state depends on its current state at time $t - 1$ and on the probability of transition between each state.

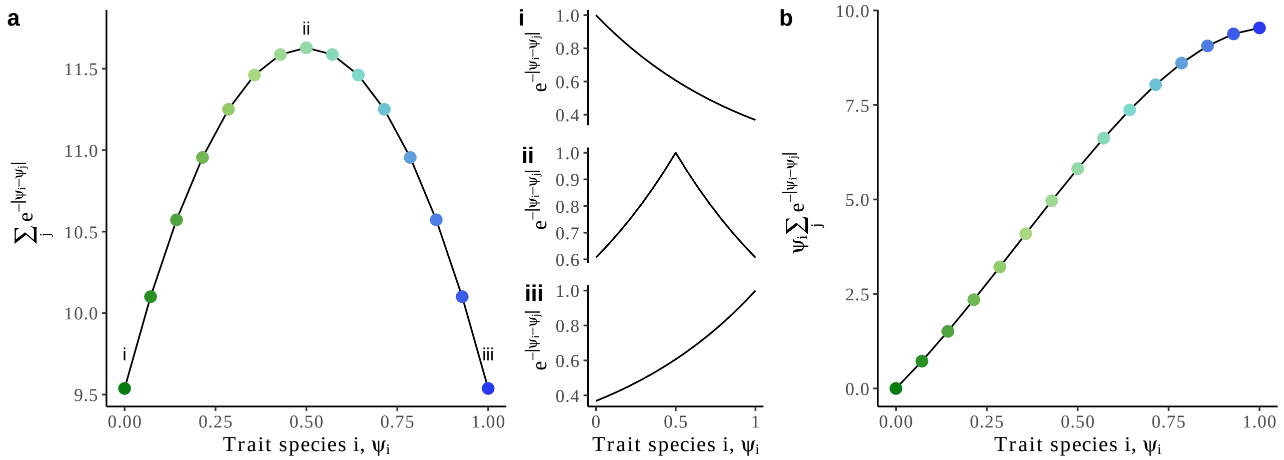


Figure S1.2: **Form of competition function favors species at the edge of the trade-off.** We show how the choice of the exponential to describe competition between the plants of different strategies, creates an advantage for species having strategies at the edge. (a) Sum of the exponential terms within the competition function. We show for 3 specific species strategies (i, ii, iii) the shape of competition function experienced by each species. (b) Similar to (a) but taking into account the effective level of competition experience (*i.e.,* with the sensitivity to competition $\psi_i$). Because of this saturation for highly stress-tolerant species (in b), it gives an advantage to the species either highly tolerant ($\psi = 1$; iii) or highly competitive ($\psi = 0$; i) as the latter doesn't suffer from the impact of strategy similarity in the competition function. This explains, why only species at the edge are present in species-rich communities under high competition. Here $N = 15$.

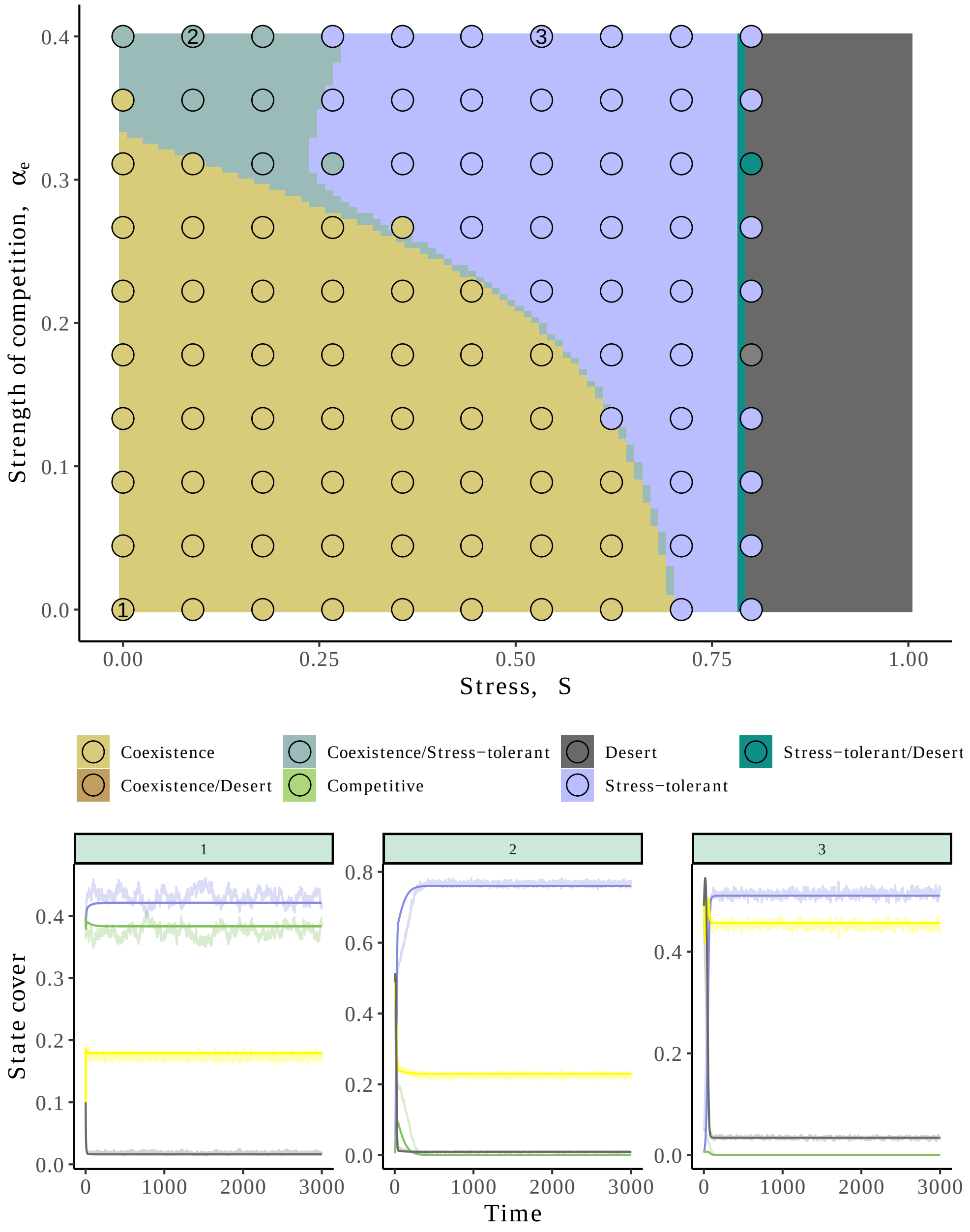


Figure S1.3: **Pair-approximation model recovers well the spatially explicit stochastic simulations.**
We compare the qualitative behavior of the pair-approximation and the model ran with cellular automata in a 100 x 100 landscape. To do so, we overlaid the result from Fig. 2a with $\delta = 0.1$ and local facilitation, and 100 simulations of the spatially explicit model equally spaced on the abiotic stress, interspecific competition space (points). While we see some deviations between the two models, potentially due to the stochasticity of the cellular automata, as well as the narrow bistability area between the vegetated and the desert states, we retrieve qualitatively the same patterns using the pair-approximation with more bistability at higher levels of competition and bistability between desert and vegetation at high abiotic stress. We show some dynamics comparing the output of the pair-approximation (deterministic) model and the spatially explicit stochastic simulations for 3 parameter sets (1, 2, 3).

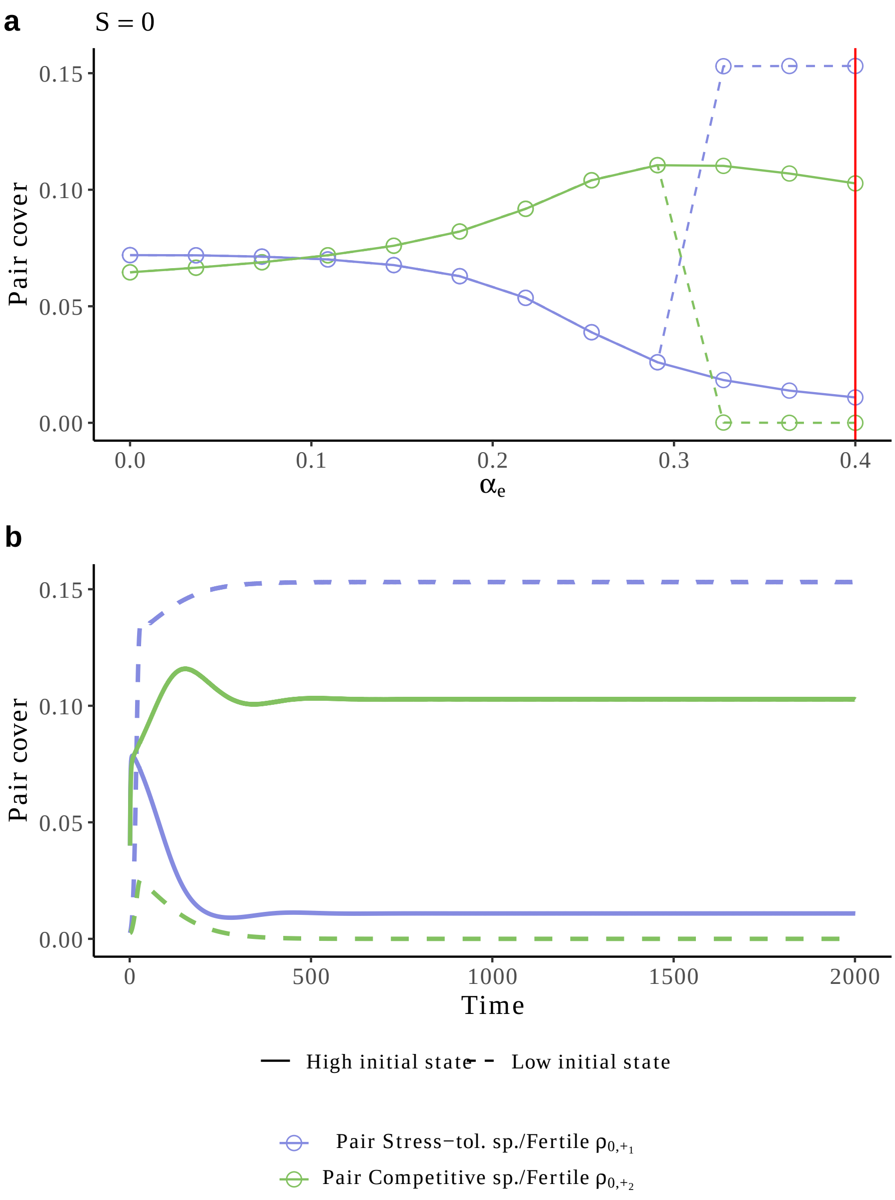


Figure S1.4: **Community bistability emerge due to the limited access of fertile sites.** (a) Changes in pair cover ($\rho_{+1,0}$ in blue, $\rho_{+2,0}$ in green) along the strength of interspecific competition. Dotted and solid lines correspond to two different initial conditions (low and high initial vegetation respectively). (b) For $\alpha_e = 0.4$ (red line in (a)), we show the temporal dynamics of pair cover for the two different initial conditions. Under high competition, when starting from an initially low vegetation state (dotted line), competition experienced by species is weak (*i.e.*, $\sum_{j=1}^{N} \rho_{+_j} \alpha_{i,j}$ is low due to low $\rho_{+_j}$). Therefore competition is not strong enough for the competitive species to exclude the stress-tolerant one that can access fertile sites in its neighborhood and rapidely dominate the landscape (positive niche construction leading to species dominance). When starting from an initially high vegetation state, both species experience more competition (but the stress-tolerant species experiences relatively more competition than the competitive one), which thereby limits its dominance and promotes species coexistence.

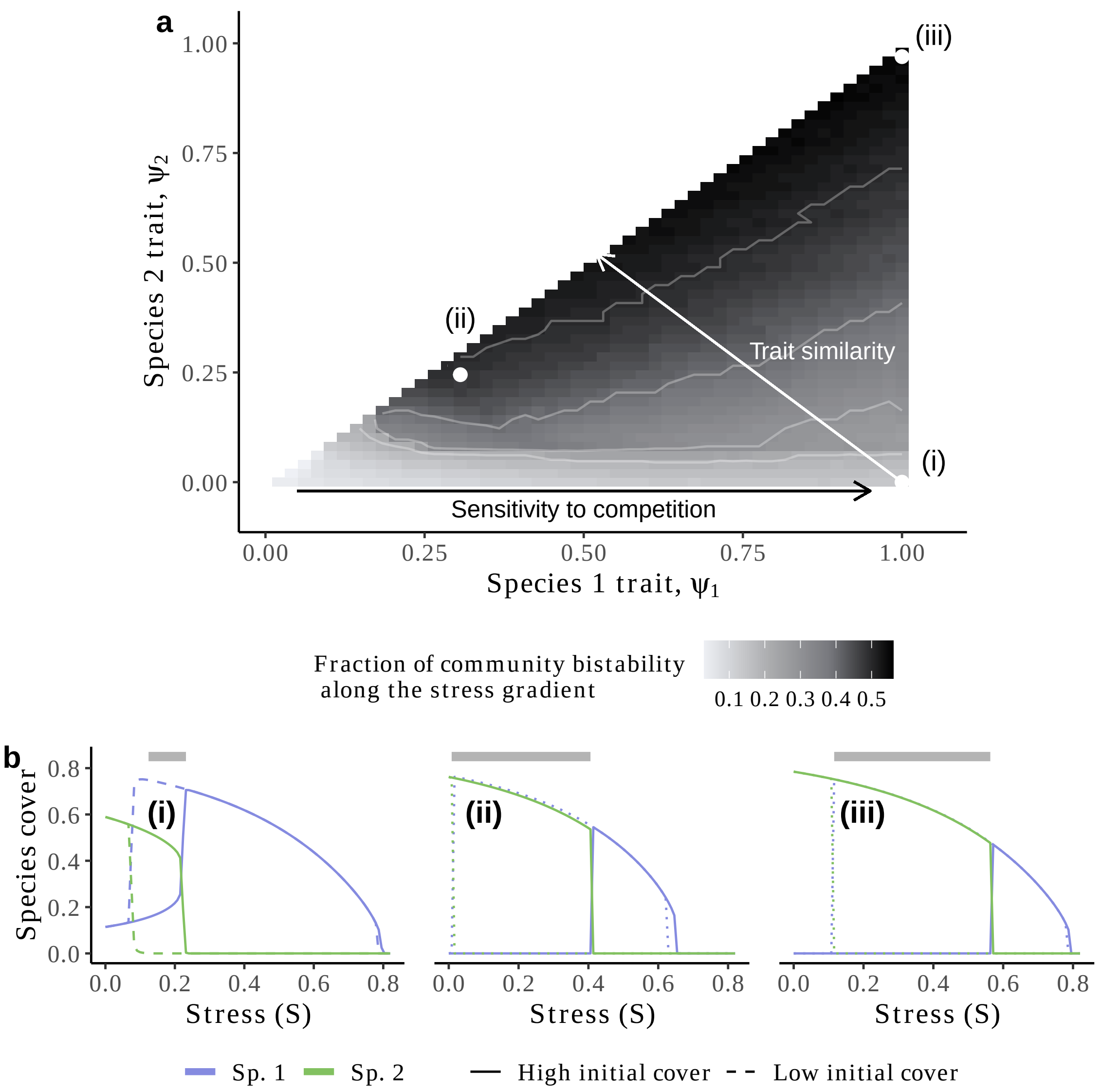


Figure S1.5: **Species relative strategies determine the occurrence of community bistability.**

(a) For each combination of species strategy ($\psi_1$, $\psi_2$), we show the fraction of the stress gradient where community bistability was found. This includes the bistability between the coexistence of species and species alone, and, each species alone in the ecosystem. To facilitate the reading, we indicate with two arrows where sensitivity to competition and strategy similarity ($|\psi_1 - \psi_2|$) between species increase. The contour lines show the isoclines for a fraction of community bistability of $0.2, 0.3, 0.4, 0.5$.

(b) For three combinations of species strategies ((i)=(1, 0), (ii)=(0.30, 0.25), (iii)=(1, 0.98)), we display the bifurcation diagrams of species cover along the gradient of stress ($S$). The linetype indicates the two trajectories of degradation (high initial cover) and restoration (low initial cover). The grey bar indicates the stress conditions over which community bistability was observed. Here, facilitation and dispersal ($\delta = 0.1$) are both local, $\alpha_e = 0.3$ and $f_0 = 0.9$. Qualitatively similar results were found for higher levels of competition (results not shown). Sp. = Species.

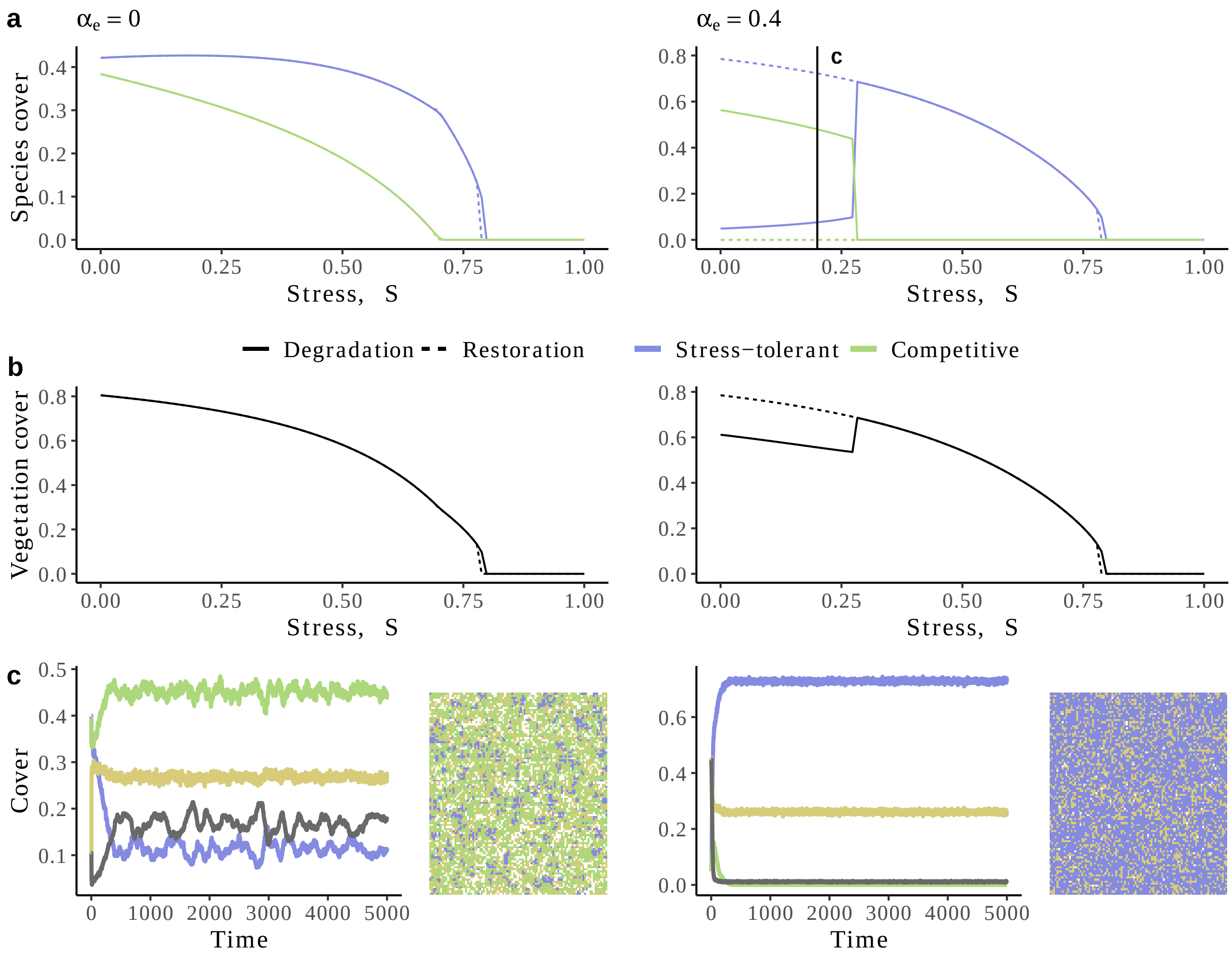


Figure S1.6: **Community succession and bistability along a gradient of abiotic stress.** The community is composed of two species, one being stress-tolerant ($\psi = 1$) and the other competitive ($\psi = 0$). (a) Bifurcation diagrams with species cover along the stress gradient ($S$) for two strengths of interspecific competition ($\alpha_e \in \{0, 0.4\}$. The color indicates the species while the linetype indicates the initial condition used (Degradation = high vegetation cover at the initial state, Restoration = low vegetation cover at initial state).
(b) Similar to (a) but with the total vegetation cover.
(c) Example time series and landscapes at asymptotic state for two different initial conditions (high and low initial biomass cover). For instance, here for $\alpha_e = 0.4$, at low stress ($S = 0.2$), depending on the initial conditions, a community with the species coexisting or the stress-tolerant species alone can be obtained. These two communities differ in their total cover ((b)-left). We show both the dynamics of the system (black line = degraded sites, yellow line = fertile soil sites) and the landscapes at asymptotic state ($t = 5000$). (a) and (b) are generated using the pair approximation model while in (c) we used the stochastic spatial simulations. Degraded sites are in white. Parameter used: $f_0 = 0.9, \delta = 0.1$ (local dispersal).

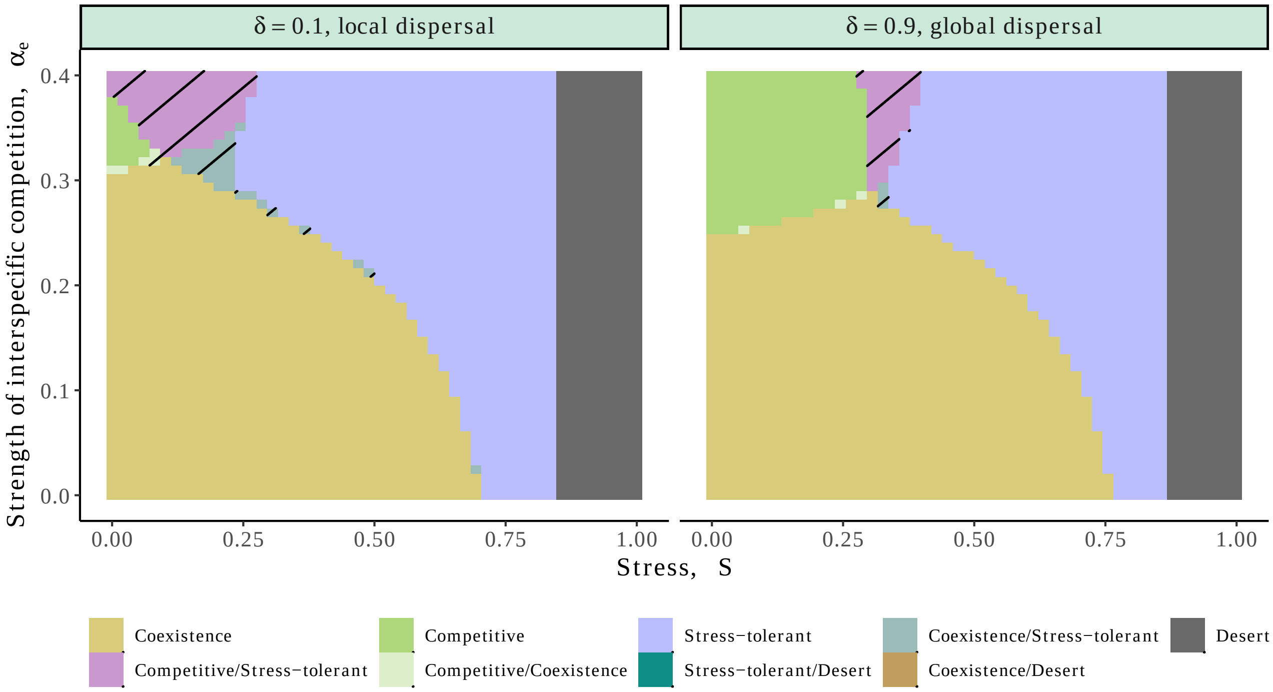


Figure S1.7: **Environmental bistability only emerges when restoration rate (*r*) can outbalance the degradation rate of fertile sites (*d*) when vegetation cover is high enough.** When basal recruitment (*r*) exceeds the degradation rate of fertile patches, competitive species do not rely anymore on the facilitation from the stress-tolerant species, and therefore, species only rarely coexist under high competition (small grey dashed area): they mainly exclude each other due to strong priority effects (pink hashed area akin to the mutual exclusion states in Fig. 5b). In addition, under these conditions, there is no bistability with the desert state: no matter the initial density of vegetation, the restoration rate exceeds the degradation rate. Here *r* and *d* were set to 0.1 and 0.05 respectively.

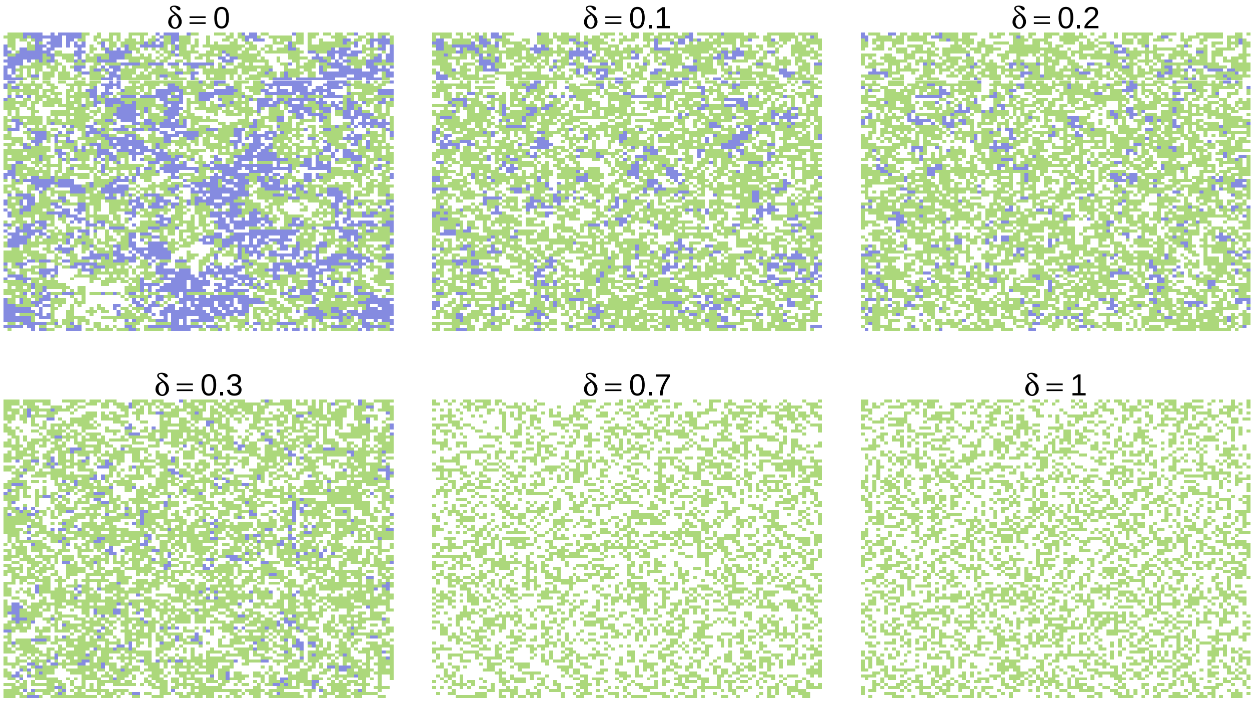


Figure S1.8: **Global dispersal benefits the competitive species and can lead to competitive exclusion of the stress-tolerant species.**
The landscape of the spatially explicit model at asymptotic state for varying dispersal rate ($\delta \in \{0, 0.1, 0.2, 0.3, 0.7, 1\}$). Here, blue = stress-tolerant species, green = competitive species, and white = degraded + fertile sites. $S = 0$ and $\alpha_e = 0.4$. Fertile and degraded sites are in white while colored cells correspond to vegetation. When dispersal is local (low $\delta$) stress-tolerant creates fertile patches where it recruits, therefore allowing both species to coexist despite the higher competitive ability for recruitment of the competitive species. Nevertheless, when seeds are spread across the whole landscape ($\delta = 1$), competitive species have more access to fertile sites and can outcompete the stress-tolerant species.

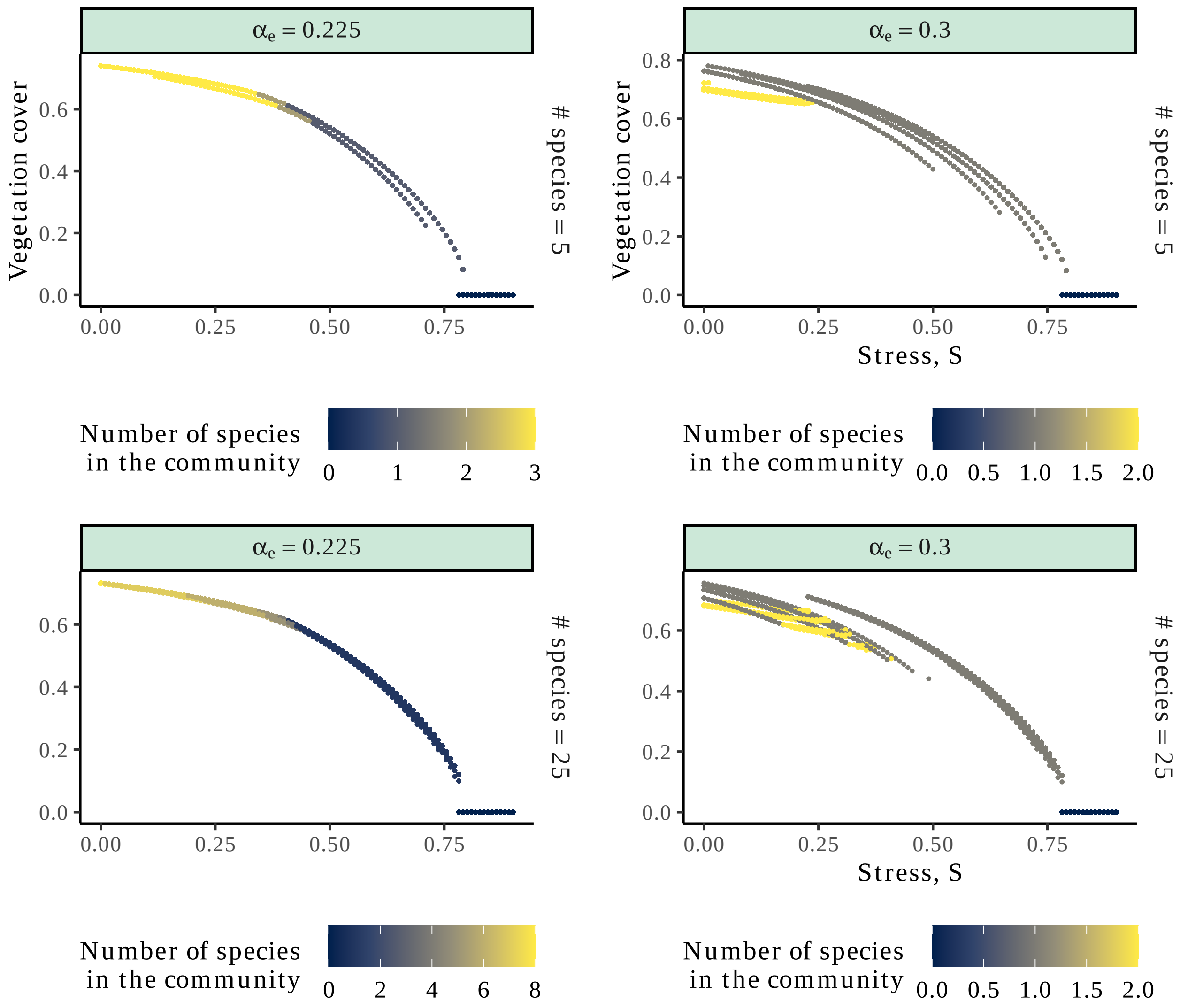


Figure S1.9: **Number of species along the stress-gradient in multispecies communities.** Same as Fig. 3 but colored with the number of species in the community.

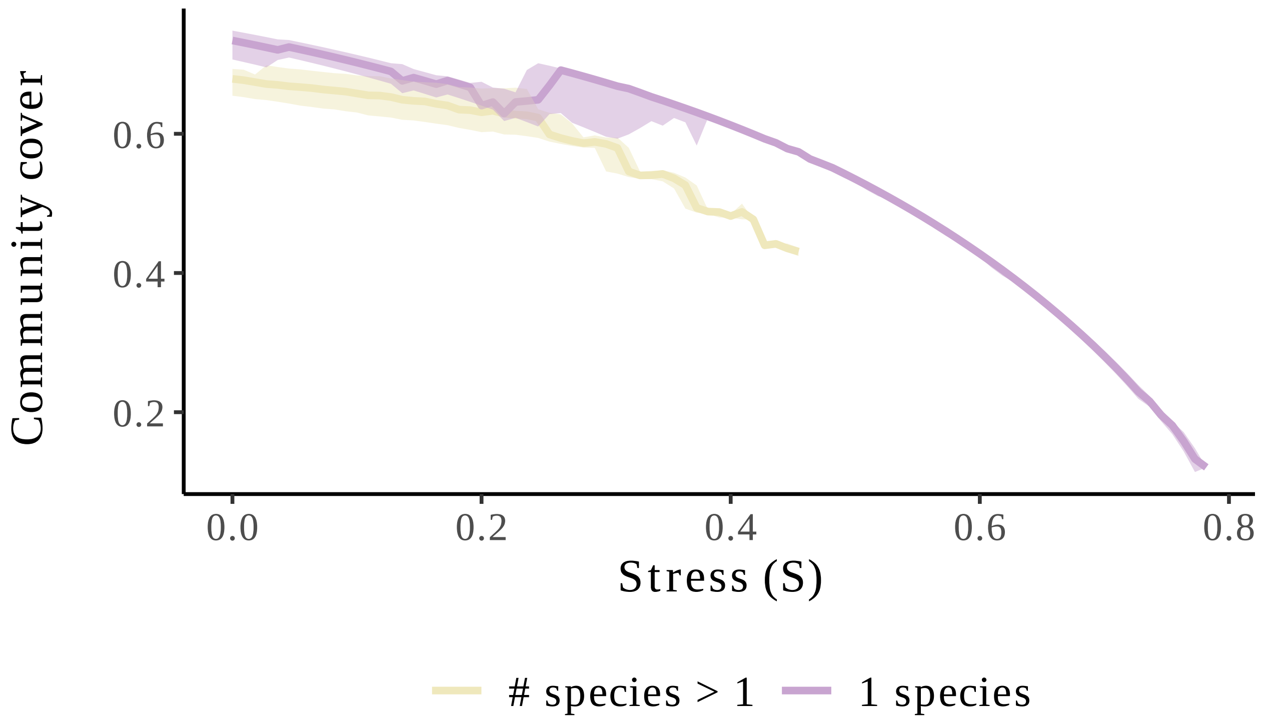


Figure S1.10: **Fingerprint of community multistability on total vegetation cover.** Total vegetation cover for alternative states of 1 (purple) or more species (yellow) averaged for all levels of competition. The median (full line) is surrounded by the first and third quartiles.

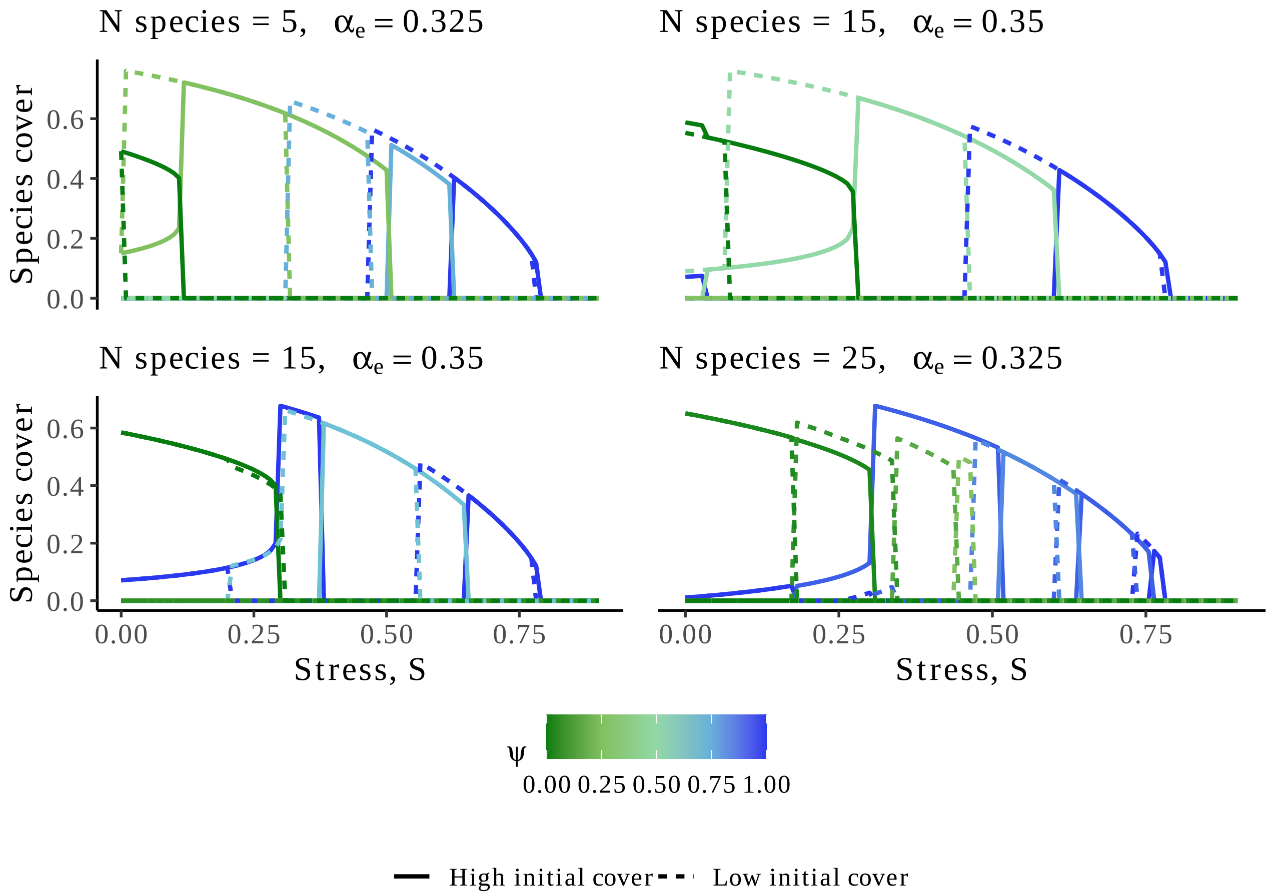


Figure S1.11: **Example of bifurcation diagrams in species-rich communities under moderate to high competition.**
We show some examples of bifurcation diagrams (for different random initial conditions) along the stress gradient for varying community richness and interspecific competition strength. The color indicates the plant strategy while the linetype indicates the type of initial condition (with high or low initial vegetation cover).

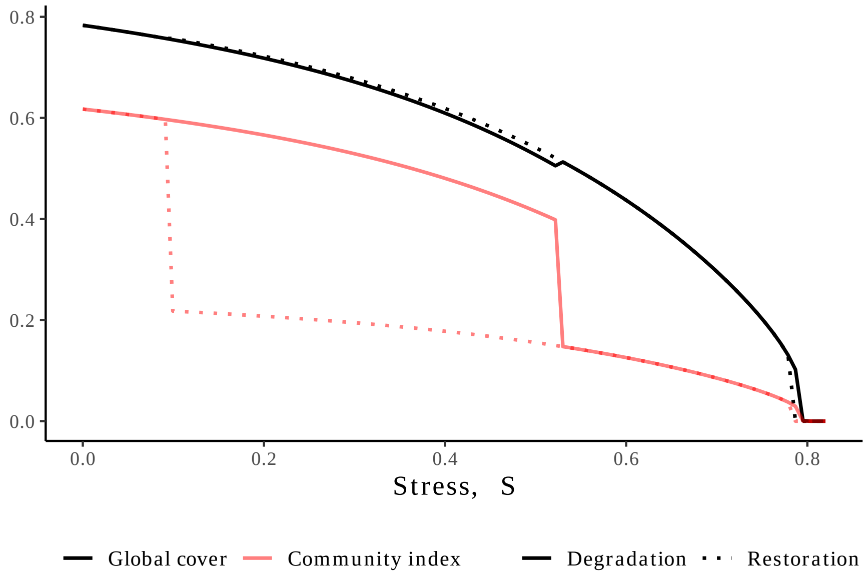


Figure S1.12: **Community index allows deciphering between alternative ecosystem states of different community composition yet with similar global vegetation cover.** Constructing bifurcation diagrams at the community level using the global vegetation cover ($\rho_+ = \sum_j \rho_{+_j}$ ; red line) does not allow to distinguish between communities of similar cover but different species composition while the community index (black line) allows such comparison along the stress gradient (S).

### S2 Relaxing the shape of the trade-off

In the main text, we assumed that the shape of the trade-off was linear. As a consequence, species at the edge of the trade-off experience less competition that the ones with intermediate strategies (Fig.S1.2). In this section, we relax this hypothesis, by exploring the behavior of the model with different shapes for the stress-tolerance/competitivity trade-off. For each term involving the strategy of a species ($\psi$), we added a power exponent $\gamma$ ($\psi^\gamma$) and explored two different shape for the trade-off: $\gamma \in \{0.5, 1.5\}$ (Fig. S2.1). Note that when $\gamma = 1$, we retrieve the linear trade-off used in the main text.

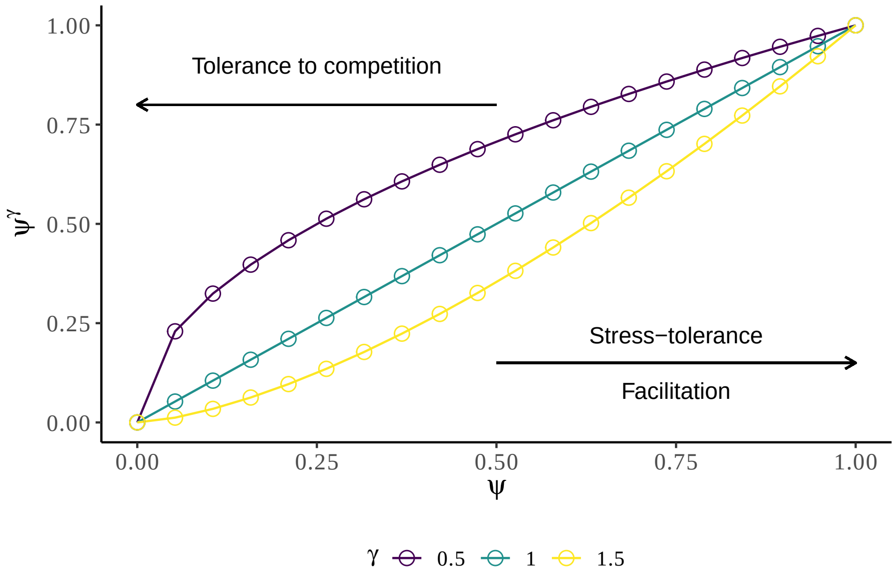


Figure S2.1: **Shape of the trade-off explored.**
Different shapes of the trade-off between competitivity and stress-tolerance are used. The case $\gamma = 1$ corresponds to the linear was used in the main text but we also explored concave ($\gamma = 1.5$) and convex ($\gamma = 0.5$) cases.

We performed the same simulations as Fig. S1.5 as the trade-off shape has no impact when species are at the edge of the trade-off: either stress-tolerant ($\psi = 1$) or competitive ($\psi = 0$) such as in Figs. S1.6, 2. In addition, we compared the degradation points and restoration points of non-linear and linear trade-offs by measuring the difference of these two thresholds between non-linear ($\gamma \in \{0.5, 1.5\}$) and linear ($\gamma = 1$) trade-offs.

We found qualitatively similar results: there is more multistability when species have more similar strategies and are more sensitive to competition (Fig.S2.2). Yet, when the trade-off is convex ($\gamma = 0.5$), even though species become more stress-tolerant and more able to facilitate their local environment, their higher sensitivity to competition increases the occurrence of community bistability. This is mainly driven by the extinction point of the least competitive species (Fig. S2.3a, b). When the trade-off shape is concave ($\gamma = 1.5$), species experience less competition and therefore, we observe more gradual changes along the stress gradient (Fig.S2.2b-(ii)). Finally, we see that it reduces the competitive advantage to the more competitive species (which persist less along the stress-gradient; Fig.S2.3b). The shape of the trade-off only slightly modulated the restoration threshold of the least competitive species (Species 2 ;Fig. S2.3c).

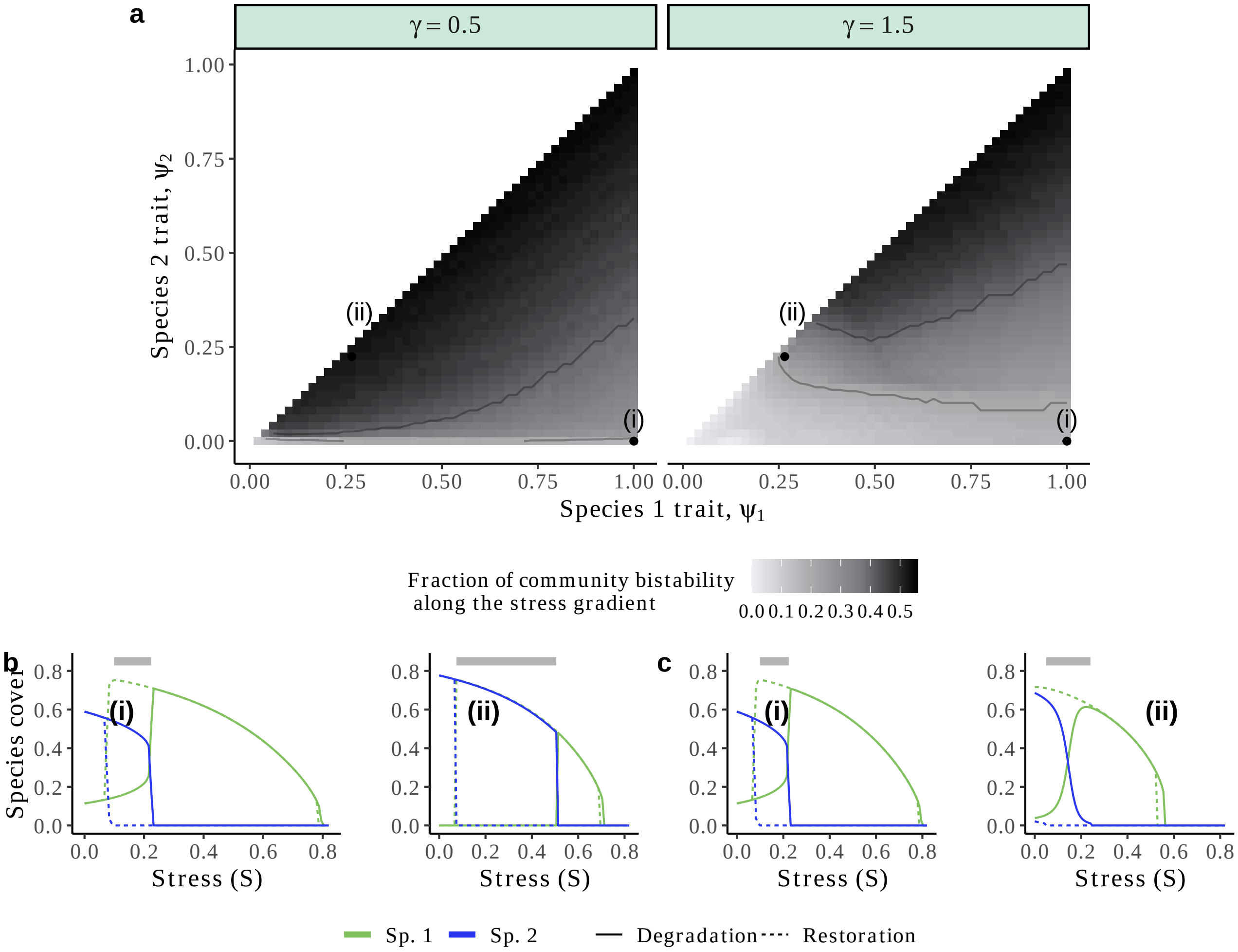


Figure S2.2: **Importance of the trade-off shape for the occurrence of community bistability in communities with varying strategies.**
(a) For each combination of species strategy ($\psi_1$, $\psi_2$), we show the fraction of the stress gradient where community bistability was found. This includes the bistability between the coexistence of species and species alone, and, each species alone in the ecosystem. The columns indicates the shape of the trade-off used ($\gamma \in \{0.5, 1.5\}$.
(b, c) For two combinations of species strategies ((i)=(1,0), (ii)=(0.27,0.22)), we display the bifurcation diagrams of species cover along the gradient of stress ($S$). The linetype indicates the two trajectories of degradation (increasing $S$) and restoration (decreasing $S$). The grey line indicates the stress conditions over which community bistability was observed. In (c), $\gamma = 0.5$ and in (d), $\gamma = 1.5$. Here, both facilitation and dispersal are local. Sp. = Species.

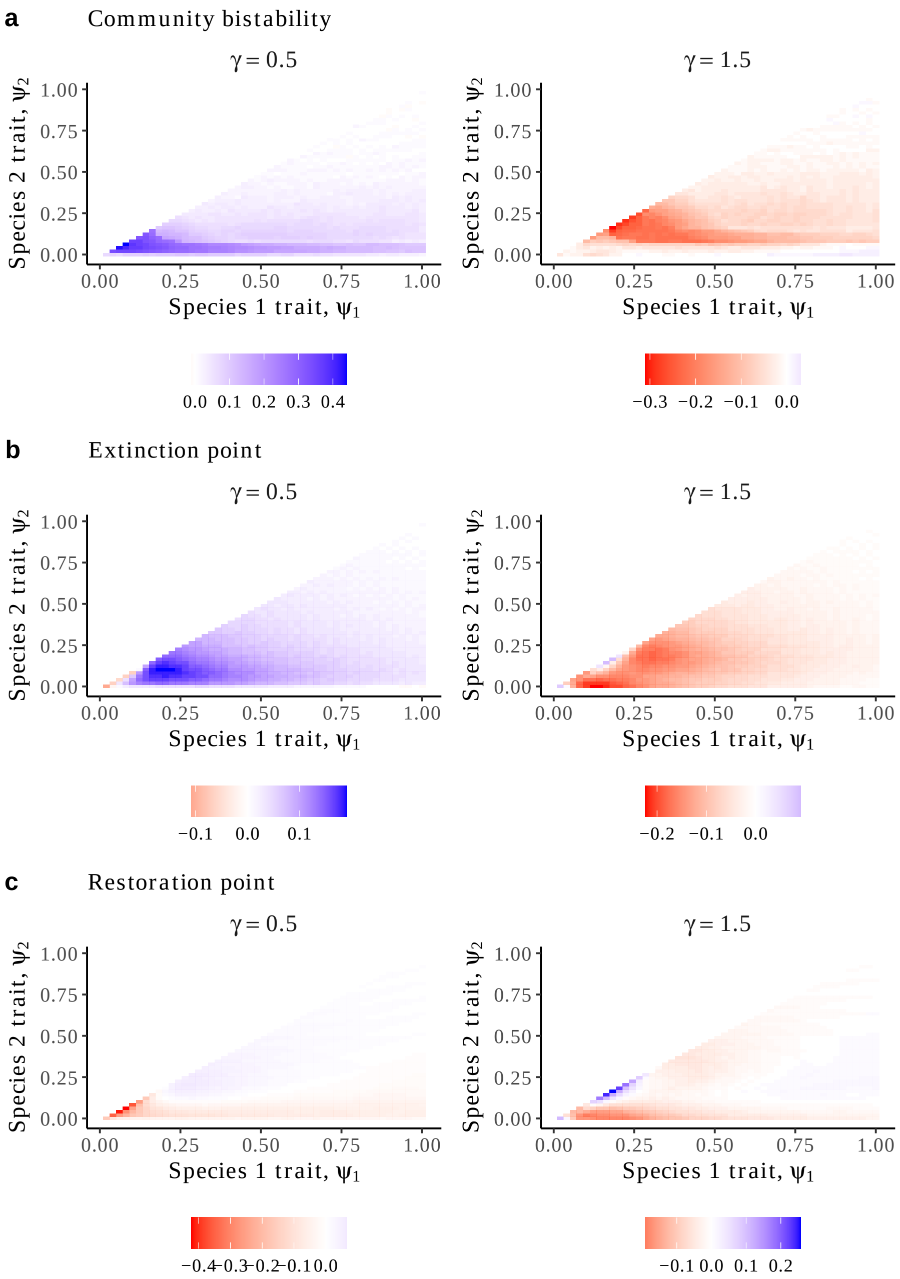


Figure S2.3: **Relative impact of the trade-off shape on the occurrence of community bistability.**
(a) Difference of fraction of community bistability between the non-linear ($\gamma \in \{0.5, 1.5\}$; columns) and the linear trade-off.
(b) Difference in the extinction point of the least competitive species (stress above which species 2 cover is 0) between the non-linear ($\gamma \in \{0.5, 1.5\}$; columns) and the linear trade-off.
(c) Difference in the restoration point of the least competitive species (stress below which species 2 cover gets positive) between the non-linear ($\gamma \in \{0.5, 1.5\}$; columns) and the linear trade-off.
In all panels, negative (*resp.* positive) values are indicated in red (*resp.* blue).

### S3 Supplementary methods

#### *Parameter values*

We used empirically derived parameters used in (Kéfi et al., 2007). Intraspecific competition strength ($\alpha_0$) was set by default to 0.3 and interspecific one ($\alpha_e$) varied between 0 and 0.4 so that interspecific competition could exceed intraspecific competition. Parameters are presented in Table. S3.1.

Table S3.1: **Details on parameters units and meaning**

| **Parameter** | **Biological meaning** | **Values** |
|---|---|---|
| $\psi$ | Species' strategy | [0,1] |
| $S$ | Stress conditions | [0,1] |
| $\delta$ | Fraction of seeds globally dispersed | [0.1, 0.9] |
| $e$ | Maximal stress tolerance | 0.1 |
| $\beta$ | Quantity of seeds produced by each plant | 1 |
| $\alpha_e$ | Interspecific competition strength | [0, 0.4] |
| $\alpha_0$ | Intraspecific coefficient strength | 0.3 |
| $m$ | Mortality rate of vegetation | 0.15 ($\sim$ 7 years) |
| $r$ | Recovery rate of degraded sites | 0.01 (100 years) |
| $f_0$ | Maximal facilitation strength | 0.9 |
| $d$ | Degradation rate of fertile soil | 0.025 (40 years) |

*Details on model assumptions*

First, we assumed that the fraction of seeds globally distributed was independent of the functional strategy of plants and the local density of vegetation ($\delta_i = \delta$; Eq. 1). Seed mass, which determines the ability of seeds to be transported by wind, herbivores, or other vectors, tends to be higher in competitive beneficiary plants compared to stress-tolerant species (Butterfield and Briggs, 2011). Therefore, we may expect competitive plants to disperse more locally than stress-tolerant ones due to enhanced local conditions near facilitating species. For instance, Valiente-Banuet et al., 2006 showed that 90% of less tolerant woody species recruited under nurses while this rate drops to 13% for nurse species. Nevertheless, Giladi et al. (2013) emphasized a higher rate of seed deposition rate of herbaceous (*resp.* shrub) species in open areas (*resp.* shrub patches) compared to shrub patches (*resp.* open areas). Therefore, we assumed a strategy-independent dispersal rate ($\delta_i = \delta$). Also, while we didn't take it into account in the model, we acknowledge that dispersal may depend on the local density of plants since shrubs in drylands can act as traps or barriers for neighboring plant seeds (Flores and Jurado, 2003; Giladi et al., 2013; Filazzola and Lortie, 2014).

Secondly, we assumed a strategy-independent mortality rate, that is $m_i = m$, where $i$ is the index of plant $i$. We acknowledge that mortality can vary between plant species in arid ecosystems (Tongway et al., 2001; Ward, 2009), in particular between shrubs and herbaceous species: while shrubs can live decades to 200 years, herbaceous species live a few years and some are annual plants. Yet, there are no clear links between plant strategies (competitive *versus* stress-tolerant) and life-expectancy (*e.g.*, Graff and Aguiar, 2017), we therefore considered strategy-independent mortality rate.

##### *Simulating the pair-approximation and mean-field model*

To build the bifurcation diagrams, we simulated the dynamics of the community until steady state for 100 values of abiotic stress ($S$) equally spaced between 0 and 1. For the two-species community, we determined the upper (*resp.* lower branch) of the bifurcation diagrams by initially setting the total vegetation cover to 0.8 (*resp.* 0.01), and the degraded and fertile sites to 0.1 (*resp.*, 0.495). Once we obtained the two branches, bistability was identified as the stress levels for which the two branches were in different states (*e.g.*, coexistence and desert state). For the N-species model, we randomly drew 250 initial species cover while keeping the total vegetation cover at 0.8 for each value of abiotic stress level.

*Definition of the community index*

In this model, alternative states can have similar vegetation cover while having different species composition. Looking just as the vegetation cover is therefore not enough to distinguish alternative states. We therefore used a community index that summarizes a system's equilibria with a single number. Following Liautaud et al. (2019), we defined this index as the scalar product of the vector of species cover at equilibrium ($\rho_{+_i}$) with a random vector $v$ of $N$ numbers between 0 and 1:

$$\text{Community index} = \sum_{i=1}^{N} \rho_{+_i} v_i$$

This random vector was fixed for all the analyses we performed. When doing the scalar product with a random vector, two different alternative states (each characterized by a unique vector of species cover) have a very low probability of being projected in the same direction, therefore allowing to distinguish communities with different species composition (see Fig. S1.12 for illustration).

Last, we counted the number of alternative stable states in the multi-species model by looking at the species composition of the alternative communities.

*Simulating the spatially explicit model*

The $2N + 2$ transitions (see below) are assumed to follow Poisson processes (*i.e.,* Markovian processes in continuous time occurring with given rates). We used a tau-leaping algorithm proposed by Gillespie (Gillespie, 2001) with a constant time step (tau-leap, $\tau = 0.5$). The idea of this method is to approximate Gillespie's algorithm on constant intervals that are sufficiently short so that the propensities of the different event types (*e.g.,* recruitment of plant $i$ summed over the landscape) do not change much during the interval. During a tau-leap ($\tau$), $2N + 2$ events can happen: recruitment of each of the $N$ plant species ($N$), their possible mortality ($N$), the degradation and restoration of fertile patches (2). Therefore, for each tau-leap:

- We compute the probabilities of each of the $2N + 2$ event at each site of the landscape and their propensity (*i.e.,* the sum of the probabilities of each of these events across the landscape)

- Then, the number of changes for each event type $j$ is drawn from a Poisson distribution of mean $\tau$ times the propensity of the event (Number event j $\sim \mathcal{P}ois(\tau \times \text{Propensity}_j)$). This gives a vector of length $2N + 2$, containing the number of transitions for each type of event across the landscape during the tau-leap

- Finally, for each event type, the sites that will be changed are drawn from a multinomial distribution with each site weighted by its local probability

- We repeat the process until an asymptotic state is reached

The simulations were run on a $100 \times 100$ landscape. For the illustrations using the spatial stochastic model, we set all species with similar initial conditions.

### S4 MEAN-FIELD MODEL

*Mean-field equations*

We analyzed the mean-field (MF) version of the model, where both the spatial structure of interactions and spatial organization of sites are not accounted for. Hence, $q_{+_i|+_j} = \rho_i$ for $i, j \in \{0, -, +_1, ..., +_N\}$ With $N$ species the equations can be described as :

$$\begin{cases} \dfrac{d\rho_{+_i}}{dt} = \rho_0 \left[ \rho_{+_i} \beta \Big( 1 - S(1 - \psi_i e) \Big) - \sum_j \rho_{+_j} \alpha_{i,j} \right] - \rho_{+_i} m \text{ for } i \in 1, 2, ..., N \\ \dfrac{d\rho_0}{dt} = \rho_- \Big( r + f_0 \sum_j \psi_j \rho_{+_j} \Big) - d\rho_0 - \rho_0 \sum_i \left[ \rho_{+_i} \left[ \beta \Big( 1 - S(1 - \psi_i e) \Big) - \sum_j \rho_{+_j} \alpha_{i,j} \right] - \rho_{+_i} m \right] \\ \dfrac{d\rho_-}{dt} = d\rho_0 - \rho_- \Big( r + f_0 \sum_j \psi_j \rho_{+_j} \Big) \end{cases}$$

*Simulations performed in mean-field*

We performed the same analysis as in the main text but with the mean-field model and found qualitatively similar patterns in terms of species succession and emergence of environmental and community bistability (Fig. S4.1). With no spatial structure, competitive species is advantaged at low-stress conditions as *(i)* it has higher competitive ability during recruitment on fertile sites compared to stress-tolerant species, and *(ii)* it benefits for both the facilitation of the stress-tolerant species (that is global facilitation for all species). Therefore, we observed less community bistability along the stress gradient compared to the pair-approximation model as the competitive species could exclude the stress-tolerant one (Fig. S4.2a), but it was still maximized when species were sensitive to competition and had similar strategies.

Finally, when compared the simulations performed with $N$ species using pair-approximation with the mean-field model and found qualitatively similar patterns (Figs. S4.3, S4.4). Interestingly, we found fewer cliques compared to the pair-approximation model suggesting that space stabilizes cliques and limits the emergence of a dominant species because the stress-tolerant species can escade competitive exclusion at low stress by facilitating its local environment and facilitating the restoration of fertile patches to recruit on (Fig. S4.5). At lower levels of stress, only mutual exclusion was observed as suggested with the 2-species mean-field model (*i.e.*, global facilitation favors most competitive species).

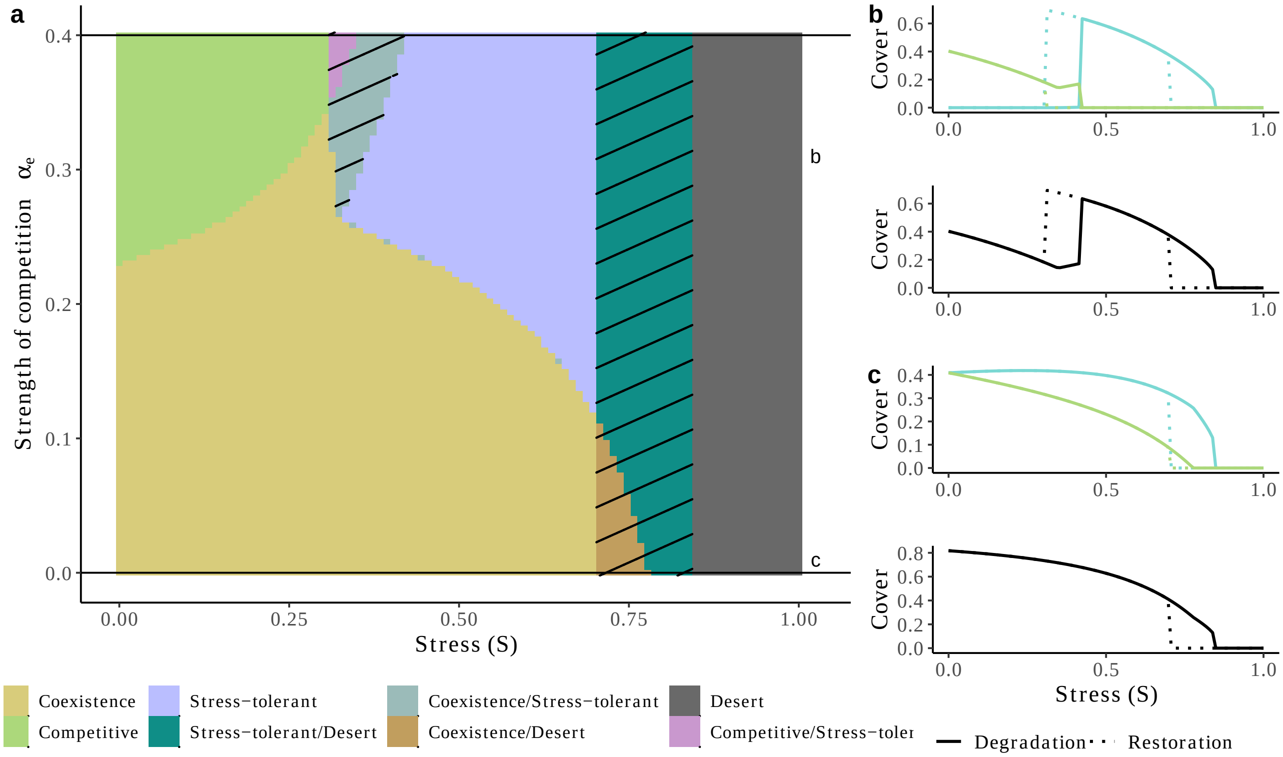


Figure S4.1: **Without spatial structure, competitive species are favored at low stress, and the occurrence of environmental bistability increases.**
The community is composed of two species, one being stress-tolerant ($\psi = 1$) and the other competitive ($\psi = 0$).
(a) Possible community states along abiotic stress gradient ($S$) and interspecific competition strength ($\alpha_e$). Areas of multistability have been hatched for better clarity.
(b) Bifurcation diagrams as the species (left; green = competitive species, blue = stress-tolerant species) and community level using the community index (right; black). The linetype indicates the two trajectories of degradation (increasing S) and restoration (decreasing S). The difference between the two trajectories indicates hysteresis. Here $\alpha_e = 0$. We also indicate the community index along the stress gradient on the top of the bifurcation diagrams.
(c) Identical to (b) with $\alpha_e = 0.4$.

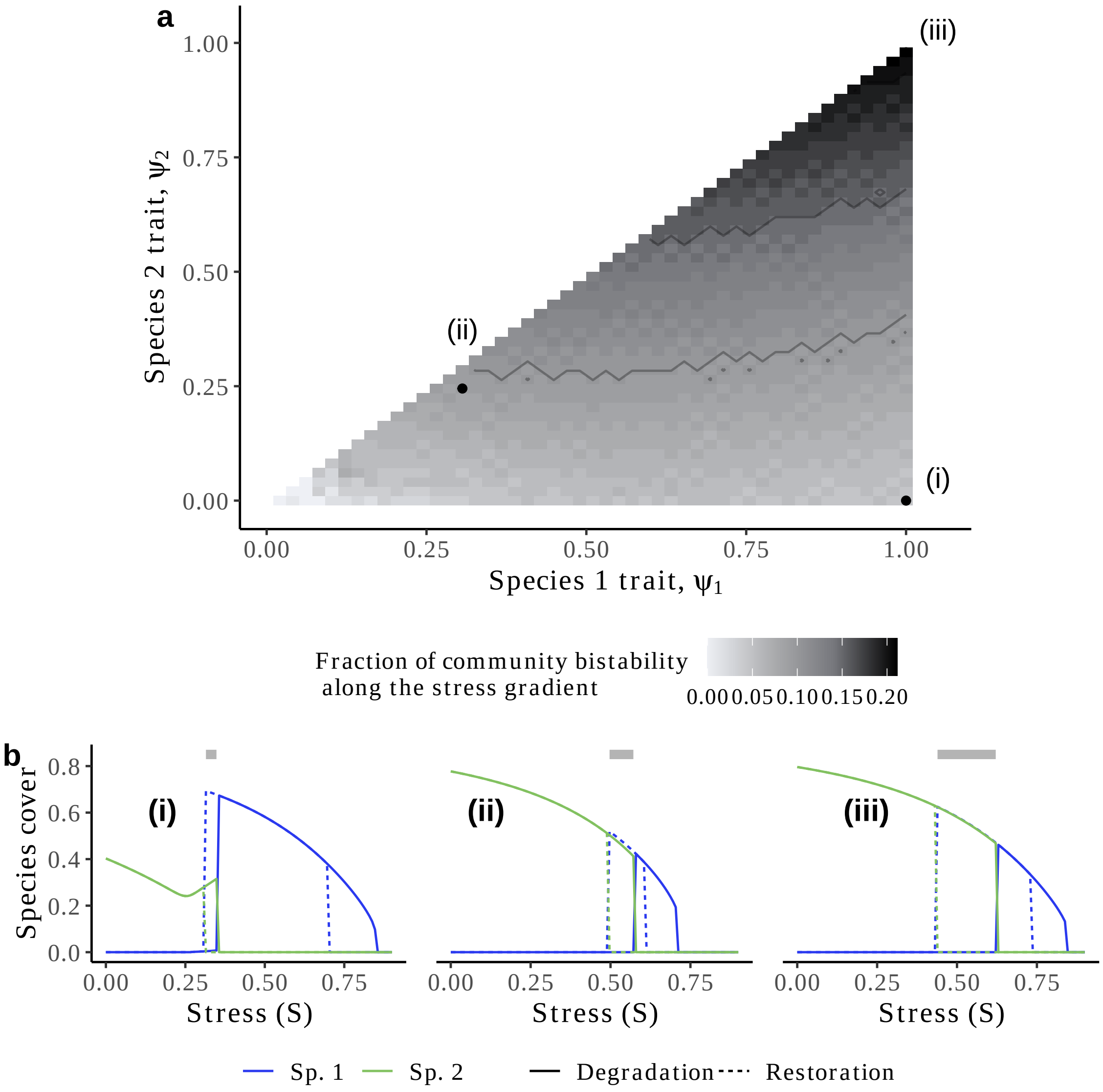


Figure S4.2: **Species relative strategies determine the occurrence of community bistability in the mean-field model.**
(a) For each combination of species strategy ($\psi_1$, $\psi_2$), we show the fraction of the stress gradient where community bistability was found. This includes the bistability between the coexistence of species and species alone, and, each species alone in the ecosystem. To facilitate the reading, we indicate with two black arrows where sensitivity to competition and strategy similarity ($|\psi_1 - \psi_2|$) between species increase.
(b) For three combinations of species strategies ((i)=(1,0), (ii)=(0.30,0.25), (iii)=(1,0.98)), we display the bifurcation diagrams of species cover along the gradient of stress ($S$). The linetype indicates the two trajectories of degradation (increasing $S$) and restoration (decreasing $S$). The grey line indicates the stress conditions over which community bistability was observed. Qualitatively similar results were found for higher levels of competition (results not shown). Sp. = Species.

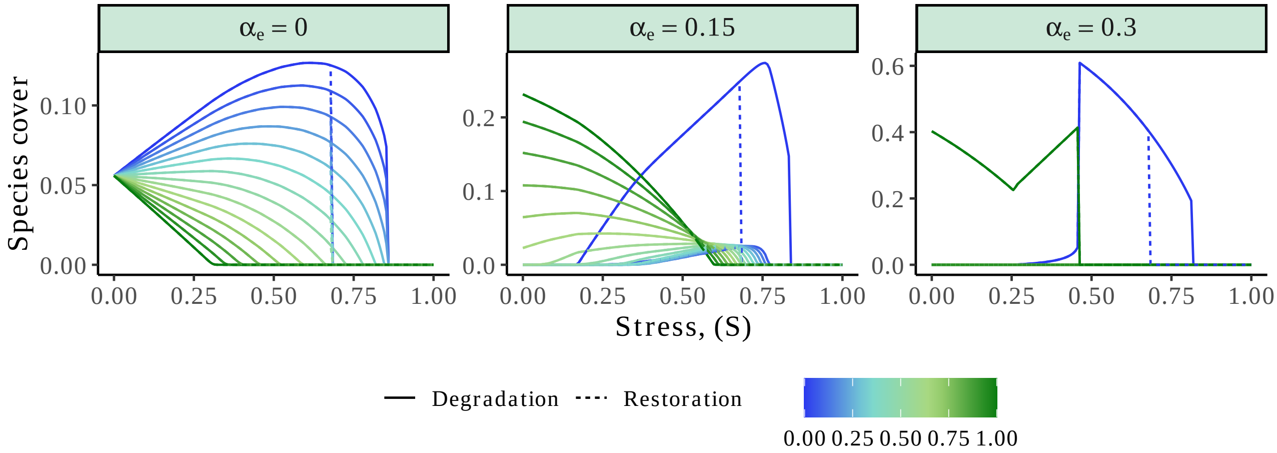


Figure S4.3: **Scaling-up to species-rich communities: the complexity of the bifurcation diagram in the mean-field model.**
(a) Bifurcation diagram along the stress gradient for $N = 15$ species. The columns indicate the strength of interspecific competition ($\alpha_e \in \{0, 0.15, 0.3\}$. The color indicates the strategy, $\psi$, of the species (from more competitive species in green to fully stress-tolerant species in blue).

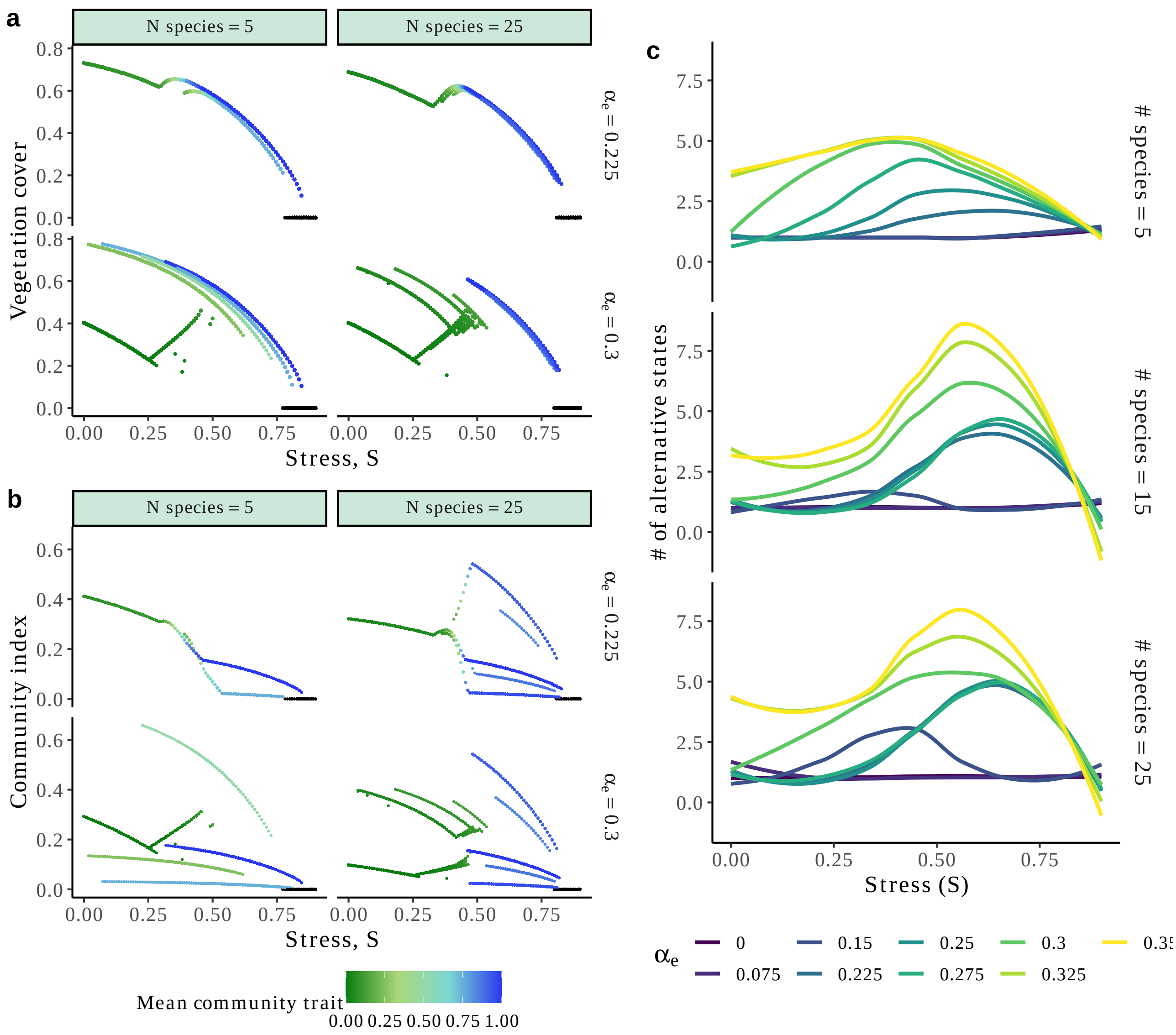


Figure S4.4: **Number of alternative stable states peaks at intermediate stress and high competition in species-rich plant communities.**
The caption is on the next page.

*Caption for Fig. S4.4.* For 5, 15, and 25 species communities, we drew 250 random initial conditions and computed for each the bifurcation diagram along the abiotic stress gradient ($S$). (a) Total vegetation cover ($\sum_j \rho_{+j}$) along stress gradient for two species richness (rows, $N = 5, 25$) and two strength of interspecific interaction (columns, $\alpha_e = 0.225, 0.35$). The color indicates the mean strategy of the community ($\bar{\psi} = \frac{\sum_j \rho_{+j} \psi_j}{\sum_j \rho_{+j}}$).
(b) Similar to (a) but with the community index to distinguish communities with similar cover but different species composition.
(c) Number of alternative stable states along the stress gradient for varying competition strength and community richness. In each panel of community richness (rows), and for each of the interspecific competition strength ($\alpha_e \in \{0.225, 0.25, 0.275, 0.3, 0.325, 0.35\}$ (colors), we counted the number of alternative stable states for each stress level as the number of different community index values that differed more than $10^{-2}$. Lines were smoothed using the *loess* method.

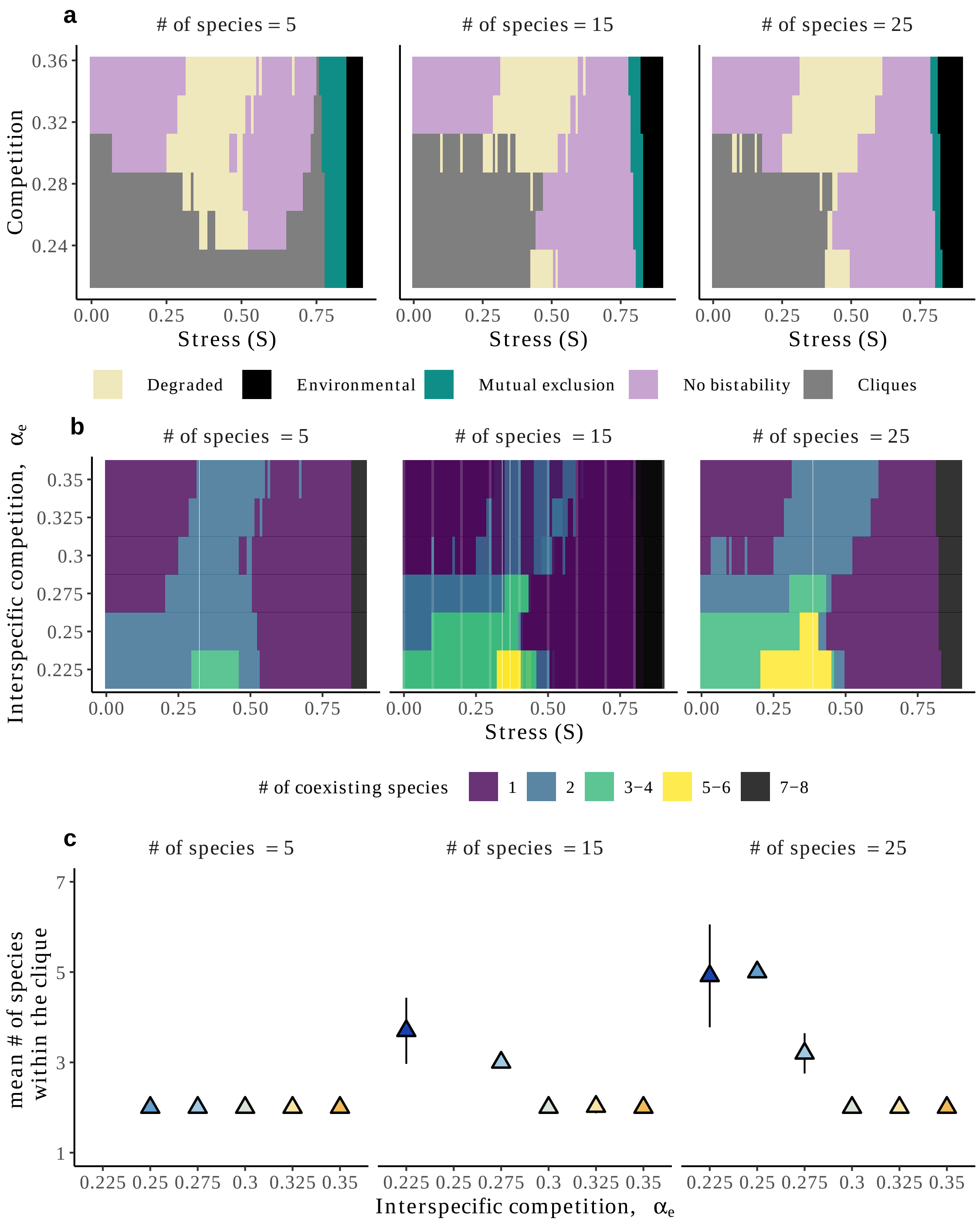


Figure S4.5: **Emergence of different types of multistability in dryland plant communities.**
(a) Types of multistability observed in plant communities of varying numbers of species (columns).
(b) Mean number of species coexisting across the 250 different initial conditions.
(c) Mean number of species within the cliques against the level of competition between species.

### S5 PAIR APPROXIMATION MODEL

In this section, we describe in more detail the pair-approximation method and the equations used for simulations. This method tracks the global densities of the $N+2$ state ($\rho_i, i \in \{-1, 0, 1, \ldots, N\}$) and the pair densities (*e.g.*, $\rho_{i,j}$). This approach supposes that the local interactions of sites can be understood by following pairs instead of triplets or quadruplets of cells (see van Baalen, 2000; Dieckmann et al., 2000 for more details). We detail the model in the case of two species and display the more general case for $N$ species.

*2 species model*

With 2 species, there is 4 global densities ($\rho_{+_1}, \rho_{+_2}, \rho_0, \rho_-$) and 10 densities of pairs as $\rho_{i,j} = \rho_{j,i}$, which makes 14 equations to completely describe the system. Yet, as we are in a spatial model where total cover sums to 1, there are 5 equations of conservation: $\sum_i \rho_{+_1,i} = \rho_{+_1}, \sum_i \rho_{+_2,i} = \rho_{+_2}, \sum_i \rho_{0,i} = \rho_0$ and $\sum_i \rho_{-,i} = \rho_-, \sum_i \rho_i = 1$, were $i \in \{+_1, +_2, 0, -\}$. This makes $14 - 5 = 9$ equations to describe the system. To not lose generally in the writing, we display the system for the arbitrary strategy of species $\psi_1 \in [0, 1]$ and $\psi_2 \in [0, 1]$ and show the system for global competition and local facilitation (as used in the main text).

$$
\begin{aligned}
\frac{d\rho_{+_1}}{dt} &= \rho_0\Big(\delta\rho_{+_1} + (1-\delta)\frac{\rho_{+_1,0}}{\rho_0}\Big) \\
&\times \Big[\beta\Big(1 - S(1-\psi_1 e)\Big) - (\alpha_{1,2}\rho_{+_2} + \alpha_{1,1}\rho_{+_1})\Big] \\
&- \rho_{+_1} m \\
\frac{d\rho_{+_2}}{dt} &= \rho_0\Big(\delta\rho_{+_2} + (1-\delta)\frac{\rho_{+_2,0}}{\rho_0}\Big) \\
&\times \Big[\beta\Big(1 - S(1-\psi_2 e)\Big) - (\alpha_{2,2}\rho_{+_2} + \alpha_{2,1}\rho_{+_1})\Big] \\
&- \rho_{+_2} m \\
\frac{d\rho_{-}}{dt} &= \rho_0 d - \rho_-\Big(r + f_0(\psi_1\frac{\rho_{+_1,-}}{\rho_-} + \psi_2\frac{\rho_{+_2,-}}{\rho_-})\Big) \\
\frac{d\rho_{+_1,+_2}}{dt} &= \rho_{+_1,0}\Big(\delta\rho_{+_2} + (1-\delta)\frac{z-1}{z}\frac{\rho_{+_2,0}}{\rho_0}\Big) \\
&\times \Big[\beta\Big(1 - S(1-\psi_2 e)\Big) - (\alpha_{2,2}\rho_{+_2} + \alpha_{2,1}\rho_{+_1})\Big] \\
&+ \rho_{+_2,0}\Big(\delta\rho_{+_1} + (1-\delta)\frac{z-1}{z}\frac{\rho_{+_1,0}}{\rho_0}\Big) \\
&\times \Big[\beta\Big(1 - S(1-\psi_1 e)\Big) - (\alpha_{1,1}\rho_{+_1} + \alpha_{1,2}\rho_{+_2})\Big] \\
&- 2\rho_{+_1,+_2} m \\
\frac{d\rho_{+_1,0}}{dt} &= \rho_{00}\Big(\delta\rho_{+_1} + (1-\delta)\frac{z-1}{z}\frac{\rho_{+_1,0}}{\rho_0}\Big) \\
&\times \Big[\beta\Big(1 - S(1-\psi_1 e)\Big) - (\alpha_{1,1}\rho_{+_1} + \alpha_{1,2}\rho_{+_2})\Big] \\
&+ \rho_{+_1,-}\Big(r + \frac{f_0\psi_1}{z} + f_0\frac{z-1}{z}(\psi_1\frac{\rho_{+_1,-}}{\rho_-} + \psi_2\frac{\rho_{+_2,-}}{\rho_-})\Big) \\
&- \rho_{+_1,0} m \\
&- \rho_{+_1,0} d
\end{aligned}
$$

$$
\begin{aligned}
\frac{d\rho_{+_2,0}}{dt} &= \rho_{00}\Big(\delta\rho_{+_2} + (1-\delta)\frac{z-1}{z}\frac{\rho_{+_2,0}}{\rho_0}\Big) \\
&\times \Big[\beta\Big(1 - S(1-\psi_2 e)\Big) - (\alpha_{2,2}\rho_{+_1} + \alpha_{2,1}\rho_{+_2})\Big] \\
&+ \rho_{+_2,-}\Big(r + \frac{f_0\psi_2}{z} + f_0\frac{z-1}{z}(\psi_1\frac{\rho_{+_1,-}}{\rho_-} + \psi_2\frac{\rho_{+_2,-}}{\rho_-})\Big) \\
&- \rho_{+_2,0}m \\
&- \rho_{+_2,0}d \\
\frac{d\rho_{+_1,+_1}}{dt} &= 2\rho_{+_1,0}\Big(\delta\rho_{+_1} + (\frac{1-\delta}{z}) + (1-\delta)\frac{z-1}{z}\frac{\rho_{+_1,0}}{\rho_0}\Big) \\
&\times \Big[\beta\Big(1 - S(1-e)\Big) - (\alpha_{2,2}\rho_{+_2,0} + \alpha_{2,1}\rho_{+_1})\Big] \\
&- 2\rho_{+_1,+_1}m \\
\frac{d\rho_{+_2,+_2}}{dt} &= 2\rho_{+_2,0}\Big(\delta\rho_{+_2} + (\frac{1-\delta}{z}) + (1-\delta)\frac{z-1}{z}\frac{\rho_{+_2,0}}{\rho_0}\Big) \\
&\times \Big[\beta\Big(1 - S(1-\psi_2 e)\Big) - (\alpha_{1,1}\rho_{+_1} + \alpha_{1,2}\rho_{+_2})\Big] \\
&- 2\rho_{+_2,+_2}m \\
\frac{d\rho_{-,-}}{dt} &= 2\rho_{0-}d - 2\rho_{-,-}\Big(r + f_0\frac{z-1}{z}(\psi_1\frac{\rho_{+_1,-}}{\rho_-} + \psi_2\frac{\rho_{+_2,-}}{\rho_-})\Big)
\end{aligned}
$$

, where :

$$
(\boldsymbol{\alpha})_{i,j} = \begin{cases} \alpha_e\big(1 + \psi_i e^{-|\psi_i - \psi_j|}\big) & \text{if i} \neq j \\ \alpha_0 & \text{else} \end{cases}
$$

*N-species model*

In the more general case of $N$-species, there are $\frac{N^2+7N+10}{2}$ equations in total. For each species $i$, we define $\psi_i$ as its position along the strategy axis. Finally, we define $A$, the matrix of competition, where $(A)_{i,j} = \alpha_{i,j}$ is the coefficient of competition of plant $j$ on plant $i$. Here are the equations for local facilitation but global competition.

$$
\begin{aligned}
\frac{d\rho_{+_i}}{dt} &= \rho_0\Big(\delta\rho_{+_i} + (1-\delta)\frac{\rho_{+_i,0}}{\rho_0}\Big) \times \Big[\beta\Big(1 - S(1-\psi_i e)\Big) - \sum_j \rho_j \alpha_{i,j}\Big] - \rho_{+_i} m \\
\frac{d\rho_0}{dt} &= \rho_-(r + f_0 \sum_j \psi_j \frac{\rho_{+_j,-}}{\rho_-}) - d\rho_0 - \sum_j \frac{d\rho_{+_j}}{dt} \\
\frac{d\rho_-}{dt} &= \rho_0 d - \rho_-(r + f_0 \sum_j \psi_j \frac{\rho_{+_j,-}}{\rho_-}) \\
\frac{d\rho_{+_i,+_j}}{dt} &= \rho_{+_i,0}\Big(\delta\rho_{+_i} + (1-\delta)\frac{\rho_{+_i,0}}{\rho_0}\Big) \times \Big[\beta\Big(1 - S(1-\psi_i e)\Big) - \sum_k \rho_k \alpha_{i,k}\Big] + \\
&\quad \rho_{+_j,0}\Big(\delta\rho_{+_j} + (1-\delta)\frac{\rho_{+_j,0}}{\rho_0}\Big) \times \Big[\beta\Big(1 - S(1-\psi_j e)\Big) - \sum_k \rho_k \alpha_{j,k}\Big] - \\
&\quad 2m\rho_{+_i,+_j} \\
\frac{d\rho_{0,0}}{dt} &= 2\sum_j \rho_{+_j,0} m + 2\rho_{0,-}(r + \frac{z-1}{z} f_0 \sum_j \psi_j \frac{\rho_{+_j,-}}{\rho_-}) - \\
&\quad 2\rho_{0,0} d - 2\sum_k \rho_{0,0}\Big(\delta\rho_{+_k} + (1-\delta)\frac{\rho_{+_k,0}}{\rho_0}\Big) \times \Big[\beta\Big(1 - S(1-\psi_k e)\Big) - \sum_j \rho_j \alpha_{k,j}\Big] \\
\frac{d\rho_{-,-}}{dt} &= 2\rho_{0,-} d - 2\rho_{-,-}(r + \frac{z-1}{z} f_0 \sum_j \psi_j \frac{\rho_{+_j,-}}{\rho_-}) \\
\frac{d\rho_{+_i,-}}{dt} &= \rho_{0,+_i} d + \rho_{0,+_i}\Big(\delta\rho_{+_i} + (1-\delta)\frac{\rho_{+_i,0}}{\rho_0}\Big) \times \Big[\beta\Big(1 - S(1-\psi_i e)\Big) - \sum_k \rho_k \alpha_{i,k}\Big] - \\
&\quad \rho_{+_i,m} m - \rho_{+_i,m}(r + \frac{f_0 \psi_i}{z} + \frac{z-1}{z} f_0 \sum_j \psi_j \frac{\rho_{+_j,-}}{\rho_-}) \\
\frac{d\rho_{0,-}}{dt} &= \frac{d\rho_-}{dt} - \frac{d\rho_{-,-}}{dt} - \sum_j \frac{d\rho_{+_j,-}}{dt} \\
\frac{d\rho_{+_i,0}}{dt} &= \frac{d\rho_{+_i}}{dt} - \frac{d\rho_{+_i,-}}{dt} - \sum_j \frac{d\rho_{+_j,+_i}}{dt}
\end{aligned}
$$

### Supplementary materials references


Butterfield, B. J. and J. M. Briggs. 2011. Regeneration niche differentiates functional strategies of desert woody plant species. Oecologia, **165**:477–487. URL http://link.springer.com/10.1007/s00442-010-1741-y.

Dieckmann, U., R. Law, and J. A. J. Metz, editors. 2000. The Geometry of Ecological Interactions: Simplifying Spatial Complexity. Cambridge Studies in Adaptive Dynamics, Cambridge University Press, Cambridge. URL https://www.cambridge.org/core/books/geometry-of-ecological-interactions/78ABF74F154A2639EDA0A7E135C9B0F1.

Filazzola, A. and C. J. Lortie. 2014. A systematic review and conceptual framework for the mechanistic pathways of nurse plants. Global Ecology and Biogeography, **23**:1335–1345. URL https://onlinelibrary.wiley.com/doi/10.1111/geb.12202.

Flores, J. and E. Jurado. 2003. Are nurse-protégé interactions more common among plants from arid environments? Journal of Vegetation Science, **14**:911–916. URL https://onlinelibrary.wiley.com/doi/10.1111/j.1654-1103.2003.tb02225.x.

Giladi, I., M. Segoli, and E. D. Ungar. 2013. Shrubs and herbaceous seed flow in a semi-arid landscape: dual functioning of shrubs as trap and barrier. Journal of Ecology, **101**:97–106. URL https://onlinelibrary.wiley.com/doi/10.1111/1365-2745.12019.

Gillespie, D. T. 2001. Approximate accelerated stochastic simulation of chemically reacting systems. The Journal of Chemical Physics, **115**:1716–1733. URL http://aip.scitation.org/doi/10.1063/1.1378322.

Graff, P. and M. R. Aguiar. 2017. Do species' strategies and type of stress predict net positive effects in an arid ecosystem? Ecology, **98**:794–806. URL https://onlinelibrary.wiley.com/doi/10.1002/ecy.1703.

Kéfi, S., M. Rietkerk, M. van Baalen, and M. Loreau. 2007. Local facilitation, bistability and transitions in arid ecosystems. Theoretical Population Biology, **71**:367–379. URL https://linkinghub.elsevier.com/retrieve/pii/S0040580906001250.

Liautaud, K., E. H. van Nes, M. Barbier, M. Scheffer, and M. Loreau. 2019. Superorganisms or loose collections of species? A unifying theory of community patterns along environmental gradients. Ecology Letters, page ele.13289. URL https://onlinelibrary.wiley.com/doi/10.1111/ele.13289.

Tongway, D. J., C. Valentin, J. Seghieri, M. M. Caldwell, G. Heldmaier, O. L. Lange, H. A. Mooney, E.-D. Schulze, and U. Sommer, editors. 2001. Banded Vegetation Patterning in Arid and Semiarid Environments, volume 149 of Ecological Studies. Springer New York, New York, NY. URL http://link.springer.com/10.1007/978-1-4613-0207-0.

Valiente-Banuet, A., A. V. Rumebe, M. Verdú, and R. M. Callaway. 2006. Modern Quaternary plant lineages promote diversity through facilitation of ancient Tertiary lineages. Proceedings of the National Academy of Sciences, **103**:16812–16817. URL https://pnas.org/doi/full/10.1073/pnas.0604933103.

van Baalen, M. 2000. Pair Approximations for Different Spatial Geometries. In U. Dieckmann, R. Law, and J. A. J. Metz, editors, The Geometry of Ecological Interactions, pages 359–387. Cambridge University Press, 1 edition. URL https://www.cambridge.org/core/product/identifier/CBO9780511525537A127/type/book_part.

Ward, D. 2009. The biology of deserts. The biology of habitats series, Oxford University Press, Oxford ; New York. OCLC: 229023535.
